# Design and characterisation of mutant and wild-type huntingtin proteins produced from a toolkit of scalable eukaryotic expression systems

**DOI:** 10.1101/492215

**Authors:** Rachel J. Harding, Peter Loppnau, Suzanne Ackloo, Alexander Lemak, Ashley Hutchinson, Brittany Hunt, Alex S. Holehouse, Jolene C. Ho, Lixin Fan, Leticia Toledo-Sherman, Alma Seitova, Cheryl H. Arrowsmith

## Abstract

The gene mutated in Huntington’s disease (HD) patients encodes the 348 kDa huntingtin (HTT) protein. The pathogenic HD CAG-expansion mutation causes a polyglutamine (polyQ) tract at the N-terminus of the HTT protein to expand above a critical threshold of ~35 glutamine residues. The effect of HD mutations on HTT is not well understood, in part due to difficulties in carrying out biochemical, biophysical and structural studies of this large protein. To facilitate such studies, we have generated expression constructs for the scalable production of HTT in multiple eukaryotic expression systems. Our set of HTT expression clones comprises both N and C-terminally FLAG-tagged HTT constructs with polyQ lengths representative of the general population, HD patients, juvenile HD patients as well as the more extreme polyQ expansions used in some HD tissue and animal models. These reagents yield milligram quantities of pure recombinant HTT protein, including many of the previously mapped posttranslational modifications. We have characterised both apo and HTT-HAP40 complex samples produced using this HD resource, demonstrating that this toolkit can be used to generate physiologically meaningful complexes of HTT. We demonstrate how these resources can produce sufficient material for protein-intensive experiments such as small angle X-ray scattering (SAXS), providing biochemical insight into HTT protein structure. The work outlined in this manuscript and the tools generated, lay a foundation for further biochemical and structural work on the HTT protein and its functional interactions with other biomolecules.

## INTRODUCTION

Huntington’s disease (HD) is a devastating inherited neurodegenerative disorder which causes a range of progressive behavioural, cognitive and physical symptoms. Incidence of HD varies in different parts of the world but HD is thought to affect between 0.42 to 17.2 per 100,000 of the population (1). There are currently no disease-modifying therapies available for patients (2). HD is hallmarked by an expansion of a CAG-trinucleotide repeat tract in exon 1 of the *HTT* gene above a critical threshold of ~35 CAG triplets (3, 4), translating to a polyglutamine (polyQ) expansion in the extreme N-terminus of the huntingtin (HTT) protein. PolyQ expanded HTT is thought to be responsible for the wide-ranging biochemical dysfunction observed in HD models and patients including proteostasis network impairment (5), transcription dysregulation (6), mitochondrial toxicity (7, 8), cellular energy imbalance (9), synaptic dysfunction (10) and axonal transport impairment (8). Although it is thought that HTT is likely a scaffold protein (11, 12), the function of HTT, wildtype or polyQ expanded, is still incompletely understood.

Biochemical investigation of the role of HTT, in either the wildtype or the disease state, is often dependent on obtaining large amounts of pure HTT protein of different polyQ lengths. The HTT protein is 3144 amino acids long (assuming a polyQ stretch of 23 residues, NCBI Reference Sequence: NP_002102.4), a potentially daunting prospect for expression and purification given its size. A number of groups have published tools and methods by which full-length HTT might be expressed and purified from either insect (13–15) or mammalian cells (16). However, the tools produced and shared with the wider community are often limited by the number of different polyQ lengths, the position of an affinity tag or their tractability for large scale production for biochemical studies. To date, the published literature reporting experiments with purified HTT protein samples remains limited. Therefore, tools and detailed methods that will enable biochemical and biophysical studies of HTT by a larger number of researchers should accelerate our understanding of the function of this elusive protein.

Towards this end, we have cloned 28 HTT constructs which allow expression of HTT protein through transient transfection of mammalian cells or viral transduction of insect or mammalian cells. Constructs have either N or C-terminal FLAG tags to assist in purification and yields of wildtype and polyQ expanded HTT protein using these systems are up to one milligram per litre of suspension culture of either insect or mammalian cells. The protein samples obtained from a simple two-step protocol are highly pure (> 90 % purity) and amenable to numerous downstream analyses and assays. Our constructs permit production of HTT in complex with the HTT binding protein HAP40, as well as in its apo form and we have characterised these HTT protein samples. This includes mapping post-translational modifications (PTMs) of the proteins derived from both insect and mammalian cells, revealing similar modification motifs to those previously reported in the literature (17–19). Both apo and HTT-HAP40 complex samples are folded, as judged by reasonable thermal melting transitions of protein samples in solution. HTT and HTT-HAP40 samples were also assessed for monodispersity using size-exclusion chromatography in tandem with multi-angle light scattering (SEC-MALS). We demonstrate how using these resources to generate large amounts of purified HTT protein sample, enables protein-intensive experiments such as SAXS. We analyzed SAXS data for apo HTT samples and the HAP40 complex, providing initial insight to the complex structure in solution.

## RESULTS

### Cloning of HTT expression constructs

Ligase-independent cloning (LIC) was used to clone the full-length HTT gene into the baculovirus transfer vector pBMDEL (**Figure 1A**), for expression of proteins in insect cells as well as in mammalian cells. In addition to the sites for LIC, the vector contains a “stuffer” fragment that includes the SacB gene, allowing negative selection on 5 % (w/v) sucrose, and a truncated VSVG fragment for pseudotyping of the baculovirus. As previously described for other HTT clones, a 15 bp repetitive element containing a mix of CAG and CAA codons was used to encode the polyQ expansion, in an effort to help maintain stability and integrity of the DNA sequence through various generations of vector propagation (20).

**Figure 1.**
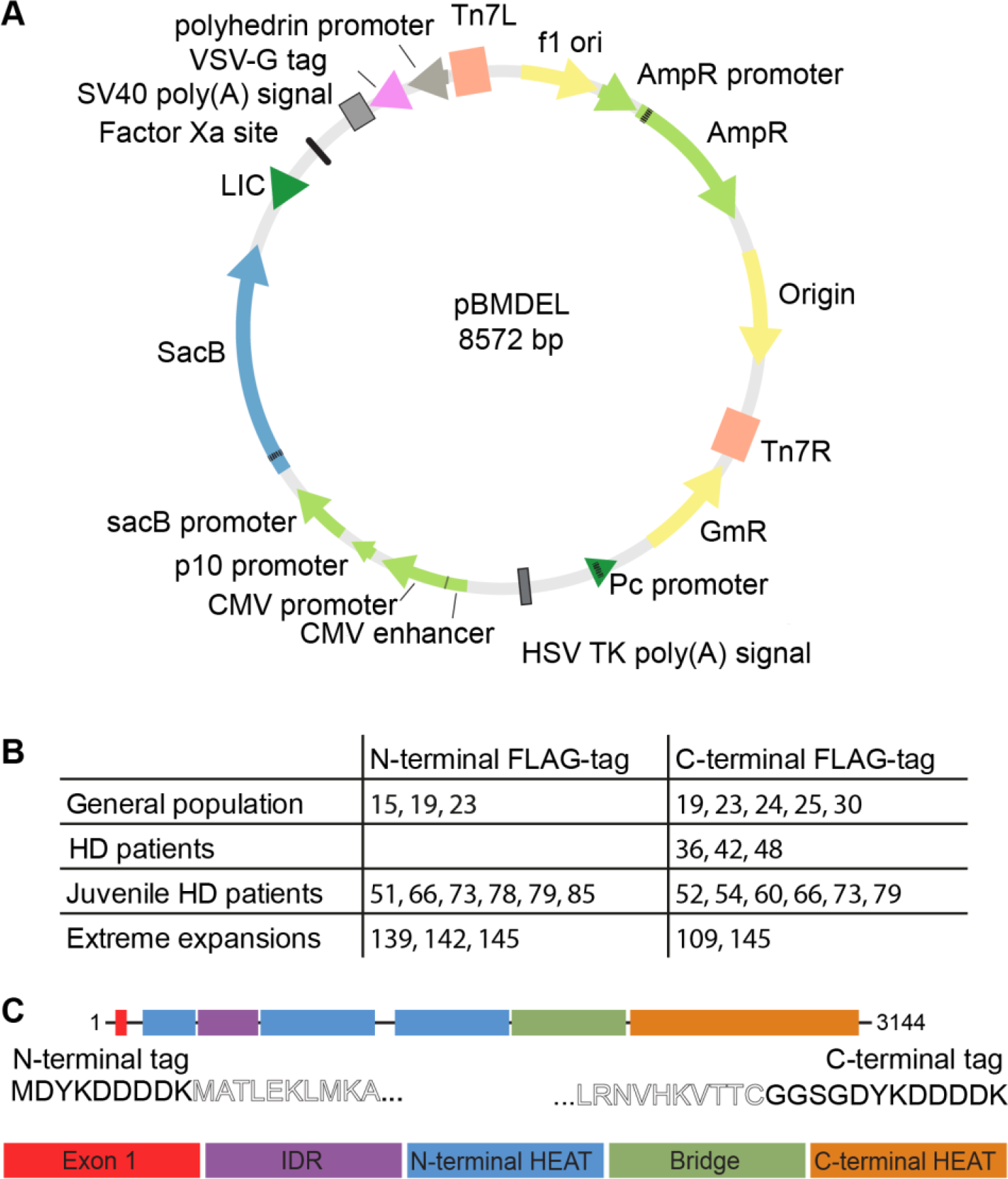
**A**) pBMDEL vector map **B**) 28 HTT expression constructs with different polyQ lengths were generated with either N or C terminal FLAG-tags, **C**) FLAG-tags are appended to either end of the full-length HTT expression construct (comprising exon1, the N-terminal HEAT domains, the intrinsically disordered region (IDR), the Bridge domain and the C-terminal HEAT domain) with minimal additional sequence

As a ~10 kb gene with multiple repetitive sequence elements, HTT is nontrivial to subclone between different vectors. We first generated N- and C-terminally FLAG-tagged pBMDEL-HTT constructs lacking part of the exon 1 sequence. Using different polyQ length encoding exon 1 PCR generated cDNAs, our LIC cloning protocol generated a variety of different polyQ encoding HTT constructs due to the error-prone nature of the recombination step. By sequencing multiple colonies we identified HTT clones with a variety of polyQ lengths with both N- and C-terminal FLAG tags (**Figure 1B**). These entry vectors serve as valuable reagents to allow future generation of even more polyQ length HTT constructs. Additionally, by using a repetitive mix of codons for the polyQ expansion (CAG CAA CAG CAA CAA)_n_, we expect improved polyQ stability over generations of plasmid, bacmid and baculovirus propagation compared to repetitive CAG codon tracts (20).

The resulting HTT open reading frames encoded within this series of constructs have either an N-terminal FLAG-octapeptide between the START methionine and the N-terminal methionine of the HTT amino acid sequence, or have the FLAG-octapeptide linked to the extreme C-terminus of HTT via a Gly-Gly-Ser-Gly linker (**Figure 1C**). As subtle changes to the exon 1 amino acid sequence of the HTT protein have been shown to give rise to changes to biophysical properties of the protein (21, 22), the C-terminally FLAG-tagged constructs allow expression of a “clean” exon 1 sequence.

### Expression of HTT variants in insect cells or mammalian cells yields functional proteins

The HTT pBMDEL expression constructs we have developed allow the expression of HTT protein by three different methods; baculovirus-mediated expression in insect cells, transient transfection in mammalian cells or transduction in mammalian cells (**Figure 2**). All three methods allow cell growth in suspension culture permitting facile scaling of the culture volumes and thus scaling of the protein production as needed. Irrespective of the expression system, HTT protein could be purified in a two-step protocol as previously described (14) from cell lysates in a Tris-salt buffer system comprising first a FLAG pull-down step and followed by size-exclusion chromatography using a Superose6 resin column (**Figure 2A**). Similar to the HTT purification efforts of other research groups, multiple peaks are present in the size-exclusion chromatography profile, likely indicating the presence of a range of different oligomeric and/or aggregated states.

**Figure 2.**
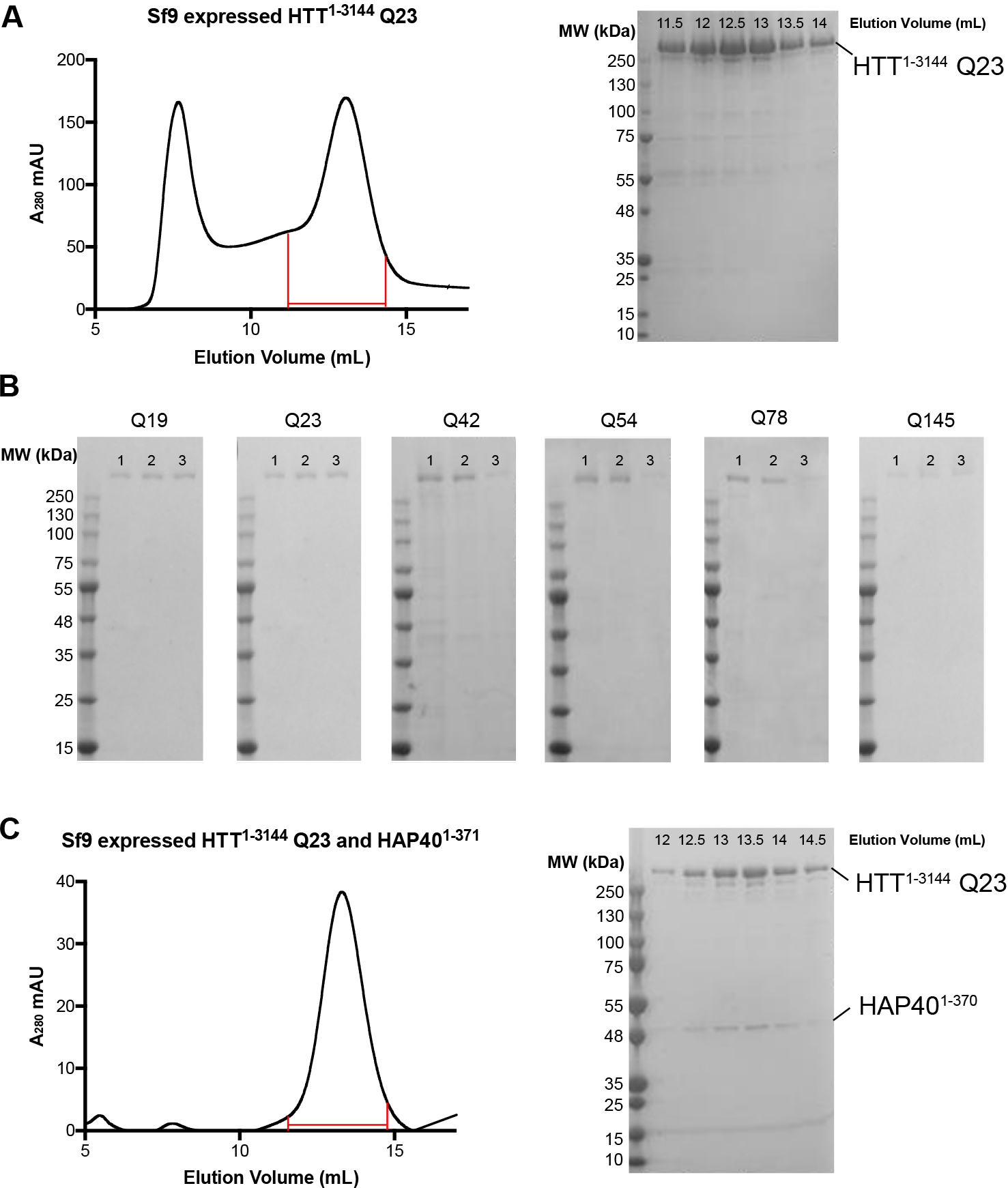
**A**) Superose6 10/300 GL column size-exclusion chromatography profile of HTT^1-3144^ Q23 expressed in Sf9 cells and purified using the C-terminal FLAG-tag. Coomassie-stained 4-20 % SDS-PAGE analysis of size-exclusion chromatography fractions (0.5 mL volume) spanning 11.5-14 mL as marked on the elution profile. **B**) Coomassie-stained 4-20 % SDS-PAGE analysis of C-terminally FLAG-tagged samples of HTT of different polyQ lengths from lane 1 – baculovirus-mediated expression in Sf9 insect cells, lane 2 - transient transfection in mammalian EXPI293F cells or lane 3 - transduction in mammalian EXPI293F cells. **C**) Superose6 10/300 GL column size-exclusion chromatography profile of HTT^1-3144^ Q23 coexpressed with HAP40^1-371^ in Sf9 cells and purified using the HTT construct C-terminal FLAG-tag and HAP40 N-terminal his-tag. Coomassie-stained 4-20 % SDS-PAGE analysis of size-exclusion chromatography fractions (0.5 mL volume) spanning 12-14.5 mL as marked on the elution profile.

Yields of the wildtype (Q23) purified HTT protein samples by the three expression methods can be as high as ~1.6 mg/L production in Sf9 insect cell culture, to ~1 mg/L in transduced EXPI293F mammalian cells and ~0.4 mg/L in transiently transfected EXPI293F mammalian cells when measured after FLAG pull-down. Comparisons of preparations of HTT Q23 samples with either N- or C-terminal FLAG tag did not show significant difference in yield. On the other hand, comparison of the yields of purified HTT with different polyQ expansions showed a trend of decreasing yield with increasing polyQ length. For example, in insect cells HTT Q42 yielded ~0.5 mg/L whereas Q145 gives yields of <0.1 mg/L. Longer polyQ lengths were also generally found to be more variable in yield between productions. For all constructs in each expression system, the 2-step purification protocol yielded a protein sample which is >90 % pure by Coomassie stained SDS-PAGE analysis (**Figure 2B**). Samples were analysed by Western blot using anti-HTT antibodies which revealed a discrete band of the expected molecular weight for each sample (**Supplementary Figure S1**).

To assess if these protein samples were folded, C-terminal FLAG-tagged HTT samples of different polyQ lengths, purified from Sf9 cells, were analysed by DSLS over a temperature gradient from 25-85 °C to assess thermal stability and propensity to aggregate under increasing temperatures. HTT samples were stable up to ~55 °C with sigmoidal thermal melting curves reflective of a folded globular protein (**Figure 3A**). Interestingly, irrespective of polyQ-length, the Temperature of aggregation (Tagg) values (35) for all HTT samples were similar at ~ 60-63 °C, indicating that the polyQ repeats did not significantly affect protein thermal stability. This suggests that polyQ may not be interacting with the folded globular part of the protein.

**Figure 3.**
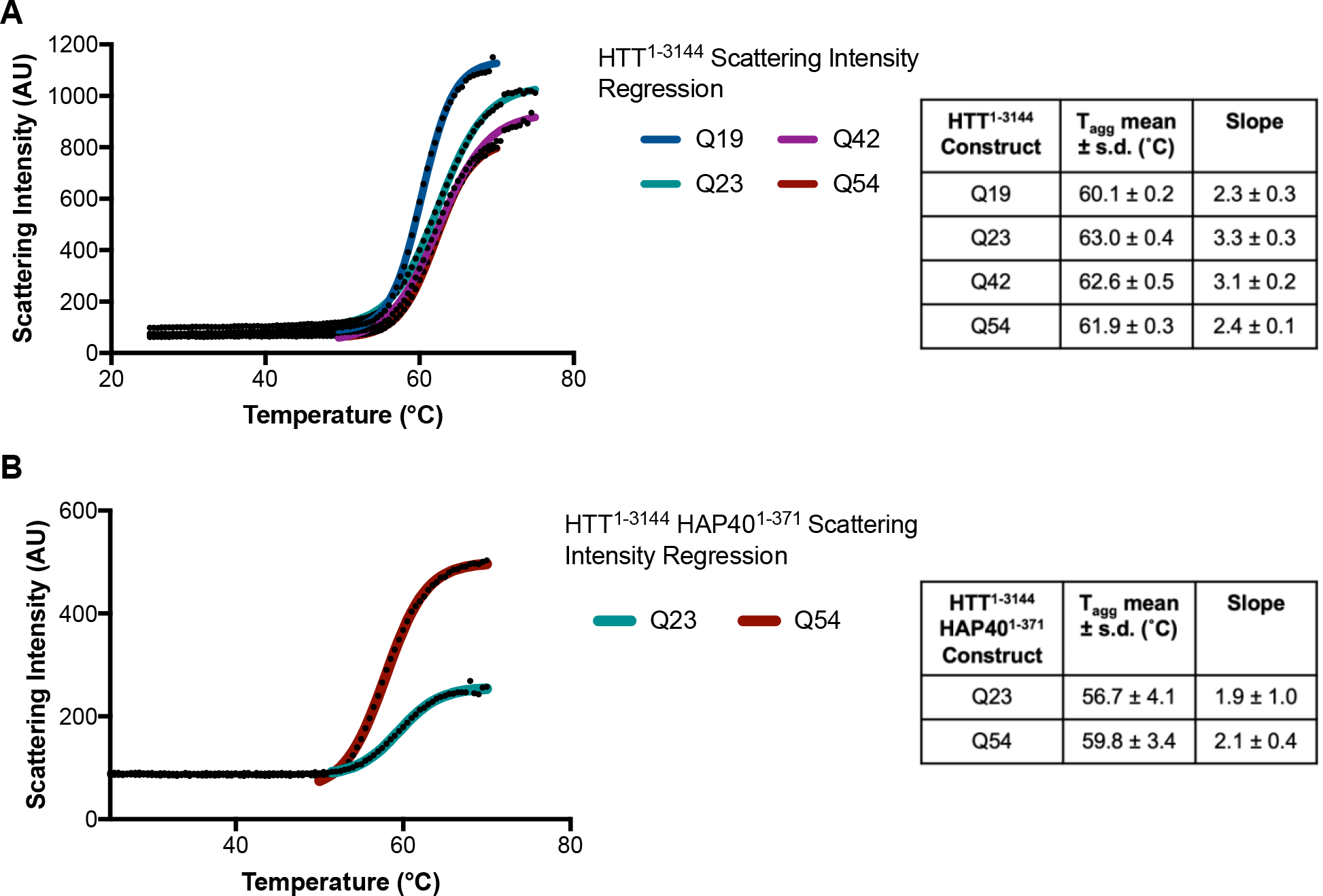
**A)** DSLS profiles of HTT^1-3144^ Q19, Q23, Q42 and Q54 (left) and the calculated thermal aggregation temperatures which show high stability indicating folded protein samples. **B)** Similar results are seen for HTT^1-3144^-HAP40^1-371^ for both Q23 and Q54 samples.

Purification of the HTT-HAP40 complex, previously reported from an adherent mammalian cell expression system (23), could be achieved through a 3:1 viral titre ratio of HTT^1-3144^ Q23 in a C-terminally FLAG-tagged pBMDEL expression construct and HAP40^1-371^ in an N-terminally His6- tagged pFBOH-MHL insect cell expression vector (**Figure 2C**). The purification protocol was modified so that an additional Ni-affinity chromatography step was included following the FLAG pull-down. The final size-exclusion chromatography step reveals that the HTT-HAP40 complex is a monodisperse sample, indicating increased protein stability and conformational homogeneity in comparison to apo HTT. Formation of this complex by HTT produced in insect cells indicates that the protein expressed is correctly folded and functional with respect to formation of an important protein interaction. HTT-HAP40 complexes with wildtype (Q23) and polyQ expanded (Q54) were also analysed by DSLS (**Figure 3B**) yielding similar thermal aggregation profiles (~57-60 °C) again, suggesting a lack of interaction between the polyQ repeat and the globular portion of the protein complex.

### HTT expressed in Sf9 insect cells retains reported phosphorylation PTMs

PTM of HTT is well described for protein derived from various mammalian cell systems and in some detail for HTT extracted from post-mortem brain tissue (17–19). However, it is not known if these PTMs are conserved in HTT expressed in Sf9 insect cells. Purified HTT Q23 and Q54 from Sf9 and EXPI293F were subjected to bottom-up proteomics (24, 25). PTMs were mapped for HTT expressed in Sf9 and EXPI293F cells and compared with published PTMs of mammalian derived HTT (**Tables 1-4** detail results for HTT Q23 samples from Sf9 and EXPI293F production, complete data can be found on PRIDE (accession PXD010865) and in Zenodo https://zenodo.org/record/2169035).

**Table 1.** Phosphorylation motifs identified for Sf9 expressed Q23 HTT^1-3144^ in reference to literature. Modifications which have been discovered in proteomics studies, but not published were retrieved from PhosphoSitePlus (17). Some modifications have not been described before. To illustrate the likelihood of these being physiologically relevant modifications, NetPhos 3.1 predictions for the putative enzyme and likelihood score are included (31). Only modifications with at least 3 peptide spectrum matches for at least one peptide containing the modification are listed in the table. All data are available via PRIDE (accession PXD010865) with summaries in Zenodo (https://zenodo.org/record/2169035). *PSMs are reported as the number of peptide spectrum matches for the most abundant peptide containing the modification described with the total number of peptide spectra for all peptides containing this modification motif in brackets.

| Site | # PSMs* | Reports or Predictions | HD Patient / control tissue samples | *In vitro* or animal models |
| --- | --- | --- | --- | --- |
| S421 | 9 (9) | Reported (16, 18, 26, 54–61, 61–79) | Yes | Yes |
| S421 / S434 | 10 (10) | Both reported | Yes/No | Yes / Yes |
| S432 | 3 (7) | Reported (16, 18, 75, 80, 81) | No | Yes |
| S434 | 3 (5) | Reported (16, 18, 54, 56, 57, 63–65, 68, 71–76, 79, 81–89) | No | Yes |
| S622 | 3 (3) | Reported (90) | No | Yes |
| S780 | 3 (4) | Predicted NetPhos 3.1 (CDK1 – 0.514) | No | No |
| T816 | 3 (3) | Predicted NetPhos 3.1 (unspecified – 0.957) | No | No |
| S1181 | 3 (5) | Reported (16, 18, 26, 57, 72, 88, 90–92) | Yes | Yes |
| S1201 | 27 (37) | Reported (16, 18, 26, 54, 55, 57, 68, 69, 74, 75, 79, 85, 86, 88–97) | No | Yes |
| S1469 | 6 (6) | Predicted NetPhos 3.1 (CDK1 – 0.513) | No | No |
| T1557 | 4 (6) | Predicted NetPhos 3.1 (unspecified – 0.777) | No | No |
| S1864 | 4 (4) | Reported (16, 18, 55, 98) | Yes | Yes |
| S1866 | 5 (8) | Reported (16, 18, 54, 55) | No | Yes |
| S1867 | 3 (5) | Predicted NetPhos 3.1 (unsp - 0.975) | No | No |
| S1876 | 4 (4) | Reported (16, 18, 54, 55, 57, 68, 70, 74, 75, 79, 83–90, 94–103) | Yes | Yes |
| Y2102 | 3 (3) | Predicted NetPhos 3.1 (unspecified – 0.921) | No | No |
| S2550 | 3 (3) | Reported (16) | No | Yes |
| S2775 | 3 (4) | Predicted NetPhos 3.1 (PKC – 0.491) | No | No |
| S2776 | 7 (8) | Predicted NetPhos 3.1 (unspecified – 0.989) | No | No |
| T2800 | 10 (13) | Predicted NetPhos 3.1 (CaM-II – 0.456) | No | No |
| S2913 | 7 (10) | Predicted NetPhos 3.1 (unspecified - 0.990) | No | No |
| T3133 | 10 (12) | Predicted NetPhos 3.1 (PKC - 0.813) | No | No |

To map the PTMs on HTT Q23 produced in Sf9 cells as completely as possible, this sample was digested with five proteases having complementary and non-specific cleavage specificity; trypsin, lysargiNase (27), pepsin, wild type α-lytic protease (WaLP) and M190A α-lytic protease (MaLP) (28). Trypsin and lysargiNase cleave at the C- and N-terminal, respectively, of lysine (K) and arginine (R) residues thus yielding complementary (or mirror-image) peptides. WaLP and MaLP preferentially cleave at aliphatic amino acids, while pepsin at pH > 2 cleaves at Phe, Tyr, Trp and Leu in position P1 or P1′ (29). MaLP, WaLP, and pepsin were selected to probe the K- and R-poor regions of the protein. Digestion of the other HTT samples was performed with trypsin alone.

When LC/MS data from the five proteolysis reactions of Q23 HTT from Sf9 are searched together, at least 90% sequence coverage was observed, while trypsin digestion of the other HTT samples yielded at least 50% sequence coverage (**Supplementary Figures S2 and S3**). As we were able to digest such large overall amounts (hundreds of micrograms) of HTT protein in multiple rounds of MS experiments, due to the high levels of production from our expression systems, this also increased the overall peptide coverage and allowed us to map PTMs with lower incidence in the samples. Due to the large size of the HTT protein, routine mass spectrometry protein identification experiments of intact sample are not feasible. Instead, we used peptide mapping analysis to confirm that the purified sample was indeed the HTT protein. As expected for the high purity of the samples indicated by SDS-PAGE analysis, HTT sequence peptides were the highest abundance proteins detected although some contaminating proteins were detected. Details of these contaminants for the HTT Q23 samples from EXPI293F cells (**Supplementary Table S1**) show that most of these proteins are unlikely to be true HTT interactors as they have high CRAPome scores (30) or are very low abundance. Modifications were detected for all samples, with well-described phosphorylation motifs being present in HTT samples from both Sf9 and EXPI293F production methods (**Figure 4**).

**Figure 4.**
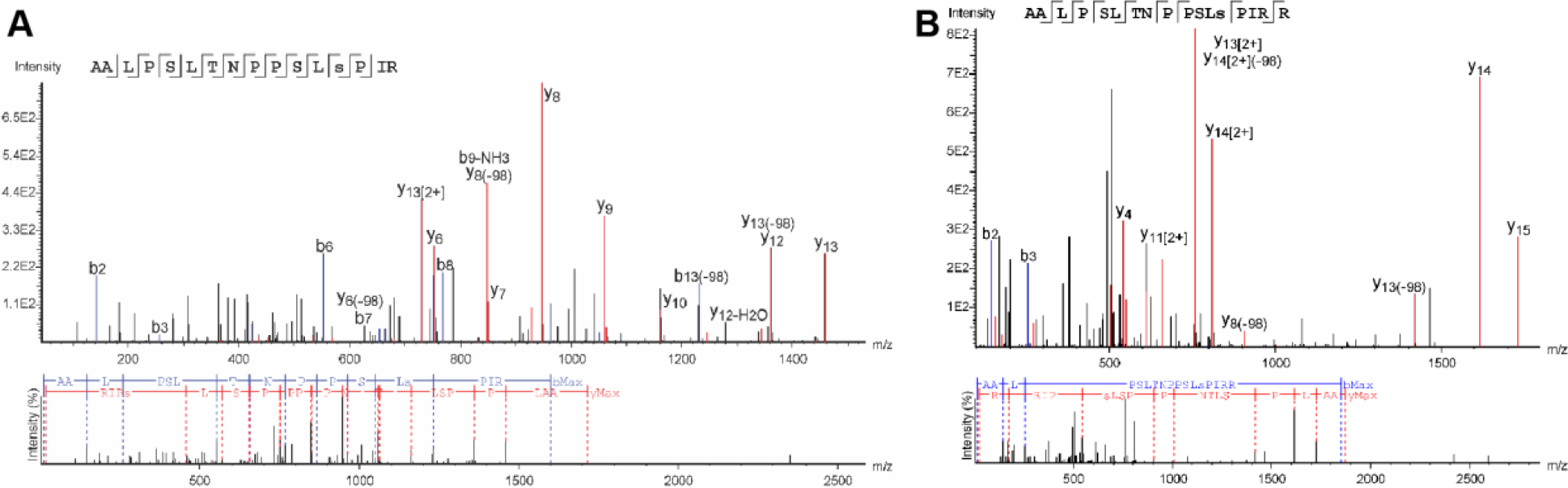
Spectra of S1181 phosphorylation motifs detected from samples of HTT^1-3144^ Q23 derived from **A**) Sf9 and **B**) EXPI293F production. This modification has been detected in human brain samples, indicating that many of the detected phosphorylation motifs detected in the purified HTT samples are physiologically relevant.

By employing multiple enzymes, sequences in regions containing sparse K and R residues, for example, a 20-amino acid long peptide within exon 1 (**Supplementary Figure S4**) were detected. Peptide-spectrum matches (PSMs) were used to prioritize the confident phosphorylation sites. HTT expressed in Sf9 cells retains many of the highly observed phosphorylation sites described in the literature for mammalian cell lines and post-mortem tissue see (**Table 1**, **Table 2** and **Supplementary Figures S5 and S6**). A total of 22 phosphorylation events with at least 3 peptide spectra matches (PSMs) were mapped in Sf9-expressed Q23 HTT, 11 of which have been reported in at least one instance in the literature. Mapping the remaining modifications to the cryo-EM HTT-HAP40 model shows that most would be surface-exposed residues in the context of apo HTT (**Supplementary Figure S7**) and their respective physiological likelihood and probably kinase, as determined by NetPhos analysis (31), is detailed in **Table 1**. Monomethylation of some lysine and arginine residues was also detected (**Table 2**). Sequence analysis of HTT using CIDER (32) and IUPred (33) in conjunction with analysis of the recently published near-atomic resolution cryo-electron microscopy structure of HTT in complex with HAP40 (23) (**Figure 5**), reveals that most of the phosphorylation sites are within disordered regions of the protein structure as described previously (18). Whilst some of these previously unreported modifications may be artefacts of the Sf9 expression system, they appear to have minimal effect on global huntingtin function, as seen by the ability of this sample to form a complex with HAP40.

**Table 2.** Arginine and lysine monomethylation modifications identified for Sf9 expressed Q23 HTT^1-3144^ in reference to literature. Methylation of huntingtin has not been previously described or reported. Only modifications with at least 3 peptide spectrum matches for at least one peptide containing the modification are listed in the table. All data are available via PRIDE (accession PXD010865) with summaries in Zenodo (https://zenodo.org/record/2169035). *PSMs are reported as the number of peptide spectrum matches for the most abundant peptide containing the modification described with the total number of peptide spectra for all peptides containing this modification motif in brackets.

| Site | # PSMs* | Site | # PSMs* | Site | # PSMs* |
| --- | --- | --- | --- | --- | --- |
| R221 | 4 (4) | R1461 | 3 (5) | K2802 | 5 (5) |
| K700 | 3 (5) | R1555 | 4 (4) | R2908 | 4 (6) |
| R908 | 7 (9) | R1947 | 3 (5) | R3112 | 3 (4) |
| K1264 | 6 (6) | K2163 | 3 (5) | R3130 | 10 (10) |
| K1448 | 3 (4) | R2774 | 7 (15) |  |  |

**Figure 5.**
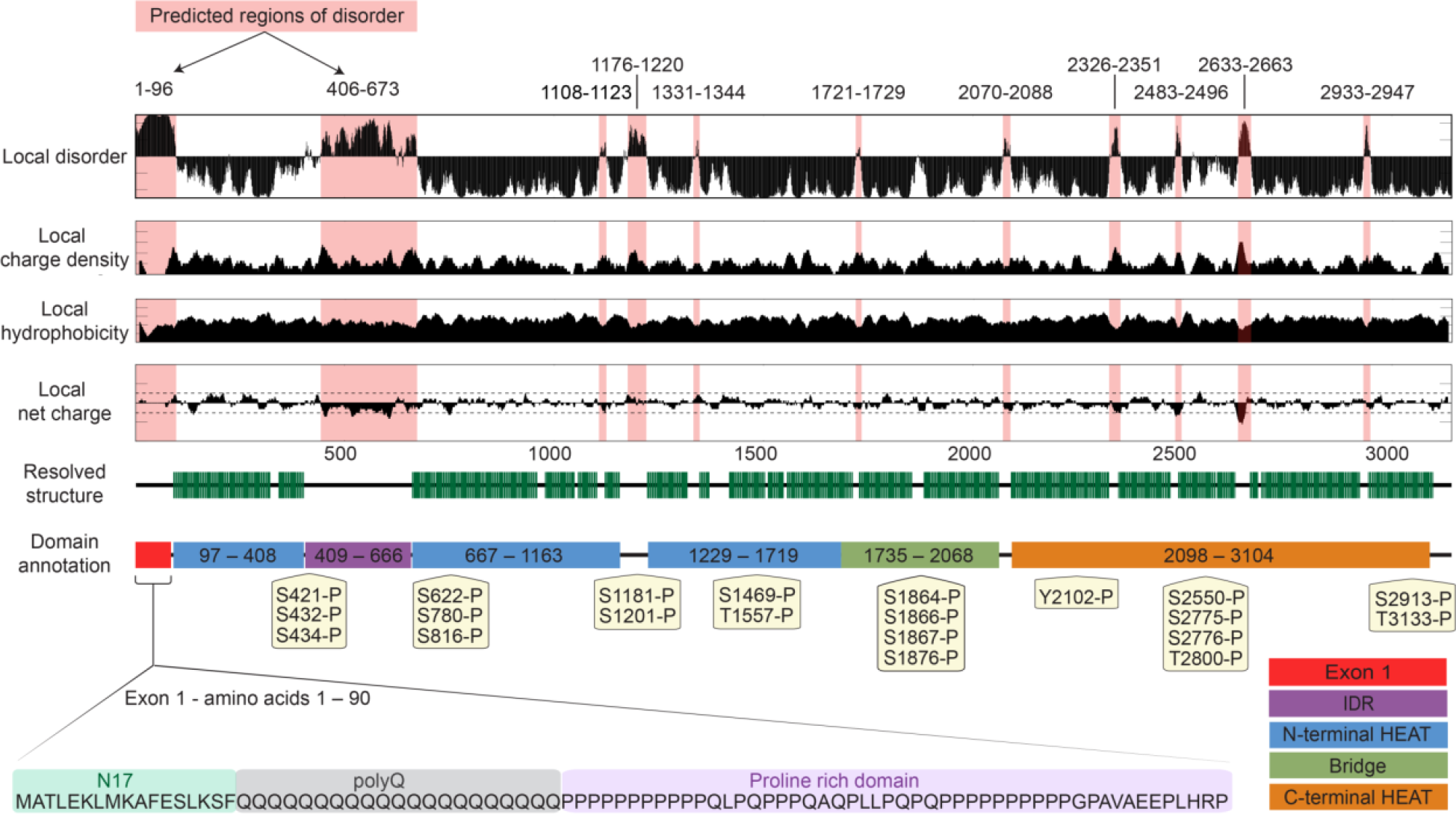
Sequence analysis of HTT complemented by structural data reveals a large modular protein connected by flexible unstructured linkers with numerous short unstructured loops protruding from the ordered domains. Local disorder, charge density, hydrophobicity and net charge are based on sequence analysis, while resolved structure and domain annotation from cryo-electron microscopy (23). Of particular note, both exon1 and the intrinsically disordered region (IDR) are not resolved in the current cryo-EM model. Phosphorylation motifs of insect-cell derived HTT Q23 mapped by peptide mass spectrometry and found in at least three peptide spectrum matches are annotated.

A total of 25 phosphorylation motifs with at least 3 PSMs were mapped for HTT Q23 expressed in EXPI293F cells, 19 of which have been described previously in the literature (**Table 3**, **Table 4** and **Supplementary Figure S5**). Interestingly, we also observed acetylation (K826 and K2932), monomethylation (R2781 and H2786) and dimethylation (R2053) of our samples, none of which have been previously described in the literature. Acetylation of HTT at other sites has been previously described and methylation motifs are observed in MS data of post-mortem human brain tissue samples of HTT (18), indicating that HTT protein methylation is a physiological modification.

**Table 3.** Phosphorylation motifs identified for EXPI293F expressed Q23 HTT^1-3144^ in reference to literature. Modifications which have been discovered in proteomics studies, but not published were retrieved from PhosphoSitePlus (17). Some modifications have not been described before. To illustrate the likelihood of these being physiologically relevant modifications, NetPhos 3.1 predictions for the putative enzyme and likelihood score are included (31). All data are available via PRIDE (accession PXD010865) with summaries in Zenodo (https://zenodo.org/record/2169035). *PSMs are reported as the number of peptide spectrum matches for the most abundant peptide containing the modification described with the total number of peptide spectra for all peptides containing this modification motif in brackets.

| Site | # PSMs* | Previously reported | HD Patient/control tissue samples | *In vitro* or animal models |
| --- | --- | --- | --- | --- |
| Y173 | 1 (1) | Predicted NetPhos 3.1 (unspecified - 0.587) | No | No |
| S419 | 1 (1) | Reported (16, 18, 60, 70, 75) | No | Yes |
| S421 | 3 (3) | Reported (16, 18, 26, 54–61, 61–79) | Yes | Yes |
| S421 / S431 | 1 (1) | Both reported | Yes / No | Yes / Yes |
| S421 / S434 | 5 (5) | Both reported | Yes / No | Yes / Yes |
| S431 | 1 (1) | Reported (16, 18, 55, 56, 63, 75, 80, 104) | No | Yes |
| S432 | 2 (4) | Reported (16, 18, 75, 80, 81) | No | Yes |
| S434 | 5 (7) | Reported (16, 18, 54, 56, 57, 63–65, 68, 71–76, 79, 81–89) | No | Yes |
| S622 | 1 (1) | Reported (90) | No | Yes |
| S623 | 3 (4) | Reported (105) | No | Yes |
| S1063 | 1 (1) | Predicted NetPhos 3.1 (unspecified – 0.989) | No | No |
| S1106 | 8 (8) | Reported (16, 90) and Zhou (2011) PhosphoSitePlus dataset (https://www.phosphosite.org/curatedInfoAction.action?record=22693604) | No | Yes |
| S1181 | 6 (8) | Reported (16, 18, 26, 54–61, 61–79) | Yes | Yes |
| S1197 | 2 (2) | Reported (16, 89) | No | Yes |
| S1201 | 12 (38) | Reported (16, 18, 26, 54, 55, 57, 68, 69, 74, 75, 79, 85, 86, 88–97) | No | Yes |
| T1262 | 1 (1) | Predicted NetPhos 3.1 (CDK1 – 0.495) | No | No |
| T1411 | 1 (1) | Reported (16, 90) and Guo (2007) PhosphoSitePlus dataset (https://www.phosphosite.org/curatedInfoAction.action?record=25582702) | No | Yes |
| T1859 | 1 (1) | Reported (106) | No | Yes |
| S1864 | 4 (5) | Reported (16, 18, 55, 98) | Yes | Yes |
| S1866 | 3 (5) | Reported (16, 18, 54, 55) | No | Yes |
| T1868 | 1 (1) | Reported (16) | No | Yes |
| S2550 | 1 (1) | Reported (16) | No | Yes |
| S2690 | 2 (2) | Predicted NetPhos 3.1 (unspecified – 0.994) | No | No |
| T2748 | 1 (1) | Predicted NetPhos 3.1 (PKC – 0.529) | No | No |
| T3098 / Y3101 | 1 (1) | Predicted NetPhos 3.1 (GSK3 – 0.446) / Predicted NetPhos 3.1 (EGFR – 0.449) | No / No | No / No |

**Table 4.** Other post-translational modifications identified for EXPI293F expressed Q23 HTT^1-3144^ in reference to literature. Methylation of huntingtin has not been previously described or reported. Only modifications with at least 3 peptide spectrum matches for at least one peptide containing the modification are listed in the table. All data are available via PRIDE (accession PXD010865) with summaries in Zenodo (https://zenodo.org/record/2169035). *PSMs are reported as the number of peptide spectrum matches for the most abundant peptide containing the modification described with the total number of peptide spectra for all peptides containing this modification motif in brackets.

| Site | # PSMs* | Modification |
| --- | --- | --- |
| K826 | 1 (1) | Acetylation (K) |
| R2053 | 1 (1) | Dimethylation (KR) |
| R2781 | 4 (4) | Methylation (KR) |
| H2786 | 6 (14) | Methylation (H) |
| K2932 | 1 (1) | Acetylation (K) |

### Characterisation of HTT and HTT-HAP40 protein sample monodispersity

Size-exclusion chromatography of HTT protein derived from insect Sf9 or mammalian EXPI293F cells using a Superose6 10/300 GL column gives a characteristic elution profile (**Figure 2A**), with a void-aggregate peak followed by peaks previously attributed as being dimer and monomer species of HTT based on column standards (14–16). The recent cryo-electron microscopy structure of HTT in complex with HAP40 reveals a bi-lobed structure of HTT in which the N-HEAT and C-HEAT domains wrap around HAP40 yielding a more compact and globular structure (23). Furthermore, HAP40 was described as being critical for producing a conformationally homogenous HTT sample amenable to cryo-electron microscopy structure determination. Our purified samples of apo HTT therefore lack a binding partner such as HAP40 which may account for the broad and overlapping elution peaks observed in gel filtration analyses as well as the tendency for self-association when HTT samples are analysed at higher protein concentrations.

To further understand this tendency for self-association and sample heterogeneity, a C-terminal FLAG-tagged HTT Q23 sample taken from the “monomer” peak of the Superose6 elution profile, was analysed by SEC-MALS using the same specification Superose6 column, which allows calculation of the in-solution protein mass. This analysis revealed a peak with a shoulder with approximate mass calculations indicating that this sample is a mixture of both HTT monomer and dimer (**Figure 6A**). In contrast, the HTT-HAP40 complex sample run on the same SEC-MALS set up at the same total protein concentration is monodisperse and the mass calculated across the peak is stable indicating the sample is homogenous and not self-associating (**Figure 6B**). Long-term storage and freeze-thaw of HTT-HAP40 samples had minimal effect on the peak profile whilst apo HTT samples had a tendency to redistribute from monomer peak to a peak profile similar to that observed during purification. Taken together, these results suggest HAP40 binding reduces homotypic HTT interaction, possibly by competing for an interaction interface or through a linkage effect (34).

**Figure 6.**
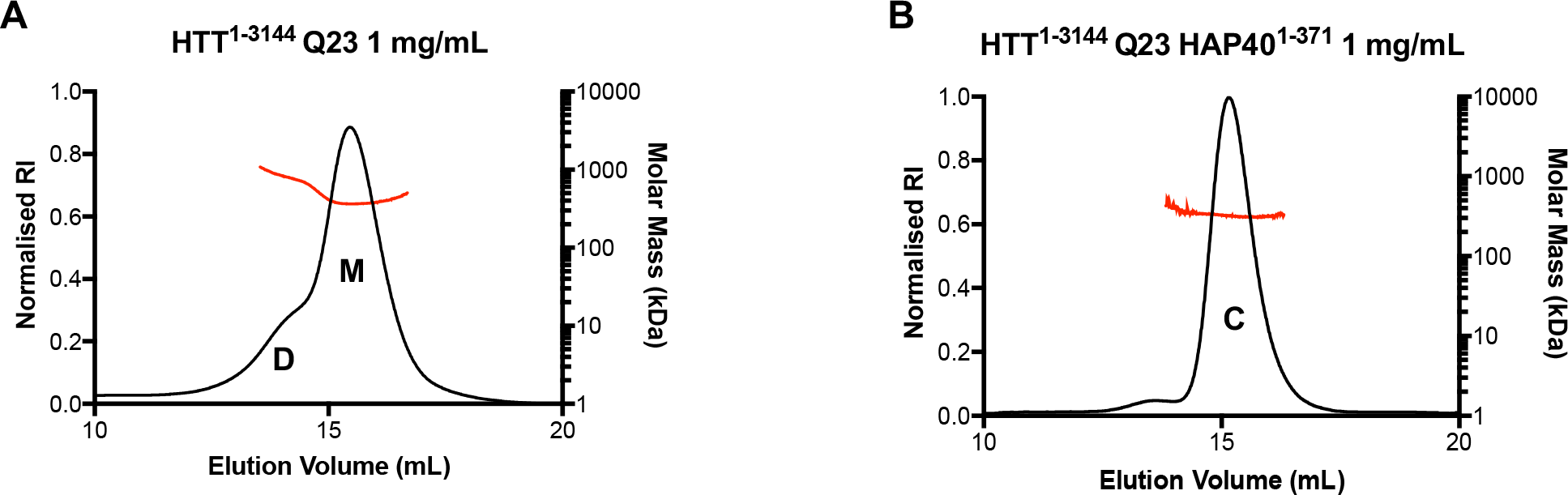
**A**) Analytical Superose6 10/300 GL gel filtration in tandem with multi-angle light scattering (SEC-MALS) profiles of HTT^1-3144^ Q23 showing distinct dimer (D) and monomer (M) species as determined by the approximate in solution mass values. **B)** SEC-MALS data for HTT^1-3144^-HAP40^1-371^ reveals that HTT-HAP40 complex (C) is more monodisperse and homogenous due to the stable mass calculated across the elution peak.

### SAXS analysis of the HTT-HAP40 complex in solution

The cryo-EM structure of HTT-HAP40 (23) has laid a tremendous foundation for our understanding of the structure of the huntingtin protein with respect to its global architecture, HEAT repeat organisation and complex formation with the HAP40 binding partner protein. However, technical limitations of cryo-EM combined with sample limitations from the conformational flexibility and heterogeneity of HTT limit our current understanding of certain structural details. The Guo *et al.* cryo-EM structure was resolved at a resolution of 4-5 Å and is missing several regions of the HTT protein. Roughly 25% of the huntingtin protein, including many functionally important elements such as exon1 and the highly modified 400-650 amino acid intrinsically disordered region (IDR), are not resolved in the cryo-EM maps, presumably due to the fact these regions are intrinsically disordered (**Figure 5**).

To further understand the structure of the entire HTT protein, we conducted Small Angle X-ray Scattering (SAXS) analysis of both the Q23 isoforms of HTT (isolated monomer peak) and HTT-HAP40 complex in solution. Similar to other biophysical and structural analysis techniques, SAXS requires large (milligram) quantities of protein. Our toolkit of HTT reagents permits production of sufficient sample for structural analyses, expediting further investigation of the HTT protein by such methodologies.

For both HTT and HTT-HAP40 sample data, Rg-based Kratky plots of the experimental curve do not fit the expected data for a generic globular protein of similar mass (**Figure 7**). The experimentally calculated radius of gyration for both samples was also much larger than that expected on the basis of the resolved residues of the cryo-EM structure (**Table 5**). This indicates that there is a degree of flexibility or disorder in both samples. This is not unexpected due to the large regions of the HTT protein sequence with predicted disorder, which are not present in the Guo *et al.* cryo-EM HTT-HAP40 model. Interestingly, the normalized pair distance distribution function P(r) of HTT-HAP40 shows a narrower range of atomic radii compared to the apo HTT sample, consistent with the HAP40 complex being more compact. However, taking into account the high propensity of HTT self-association observed in our analytical size-exclusion chromatography profiles, apo HTT SAXS analysis may also be confounded by self-association of molecules in the concentrated solutions used for SAXS data collection. This assertion is corroborated by the higher molecular weight estimated from SAXS data (**Table 5**) for apo HTT compared to HTT-HAP40. Therefore further SAXS analysis of the apo protein was not pursued.

**Table 5.**
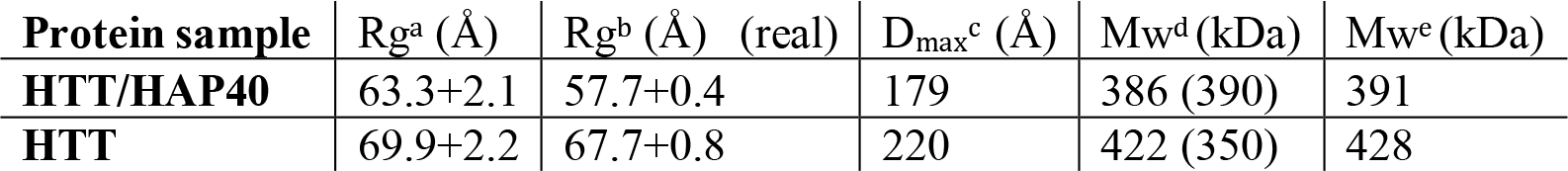
SAXS parameters for data validation and interpretation. a) Radius of gyration calculated using Guinier fit in the *q* range 0.015 < *q* < 0.025 Å^−1^ and 0.012< *q* < 0.019 Å^−1^ for HTT/HAP40 and HTT, respectively. b) Radius of gyration calculated using GNOM. c) Maximum distance between atoms from GNOM. d) Molecular weight estimated using SAXSMoW (47). MW expected from sequence is shown in the parentheses. e) Molecular weight estimated from SAXS data based on Vc (48).

| Protein sample | Rg<sup>a</sup> (Å) | Rg<sup>b</sup> (Å) (real) | D<sub>max</sub><sup>c</sup> (Å) | Mw<sup>d</sup> (kDa) | Mw<sup>e</sup> (kDa) |
| --- | --- | --- | --- | --- | --- |
| HTT/HAP40 | 63.3+2.1 | 57.7+0.4 | 179 | 386 (390) | 391 |
| HTT | 69.9+2.2 | 67.7+0.8 | 220 | 422 (350) | 428 |

**Figure 7.**
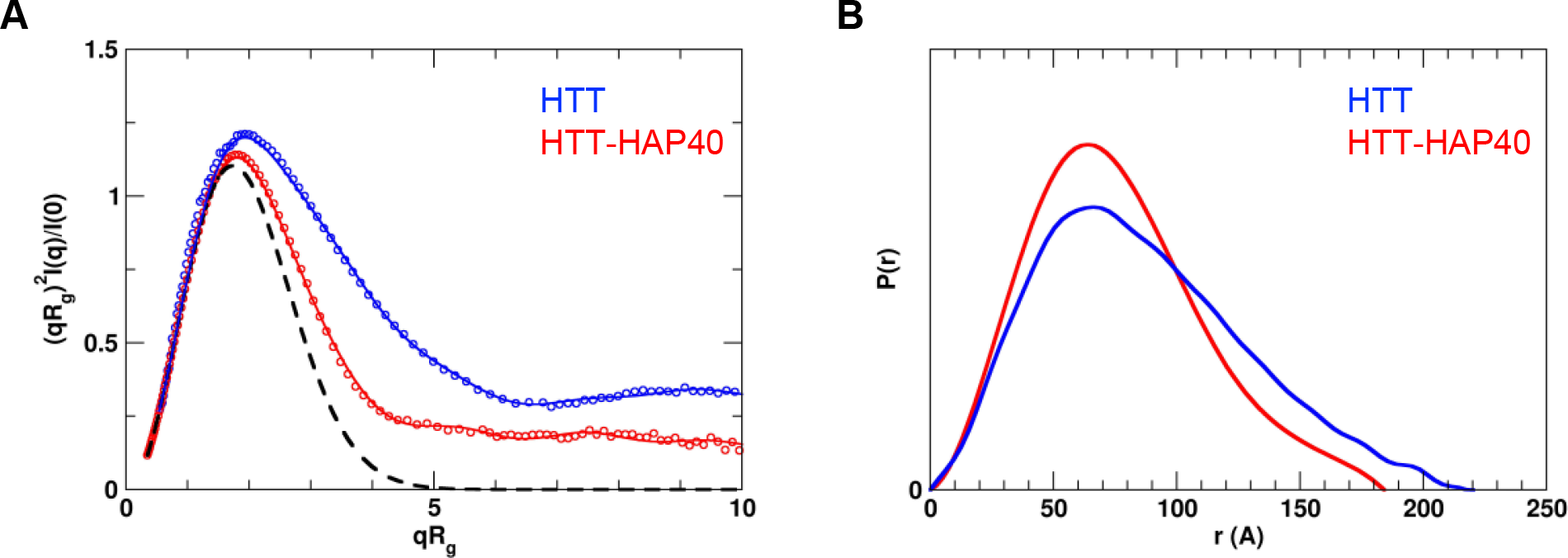
Experimental SAXS data. **A**) Rg-based Kratky plots of experimental SAXS data for HTT-HAP40 complex (red) and apo HTT (blue). The experimental data are displayed as empty circles. The solid lines show regularized experimental curves determined using GNOM. The theoretical curve expected for a spherical protein of similar size is shown by a dashed line. **B**) Normalized pair distance distribution function P(r) calculated from experimental SAXS data with GNOM.

To better understand the HTT-HAP40 structure, including the disordered/missing regions of the cryo-EM model, we performed coarse-grained molecular dynamics (MD) simulations, and calculated an ensemble of conformations that best fits the solution SAXS data for HTT-HAP40. This modelling approach assumed that the residues with known coordinates in the cryo-EM model form a quasi-rigid complex while the residues with missing coordinates are flexible. Predicted SAXS scattering curves were averaged over an ensemble of MD simulated structures using the optimal weights for each ensemble member obtained with the Sparse Ensemble Selection (SES) method (35). The resulting average “theoretical” scattering curve for the SES weighted ensemble of structures gave a much better fit to the experimental scattering data than that of the cryo-EM structure (**Figure 8A**).

**Figure 8.**
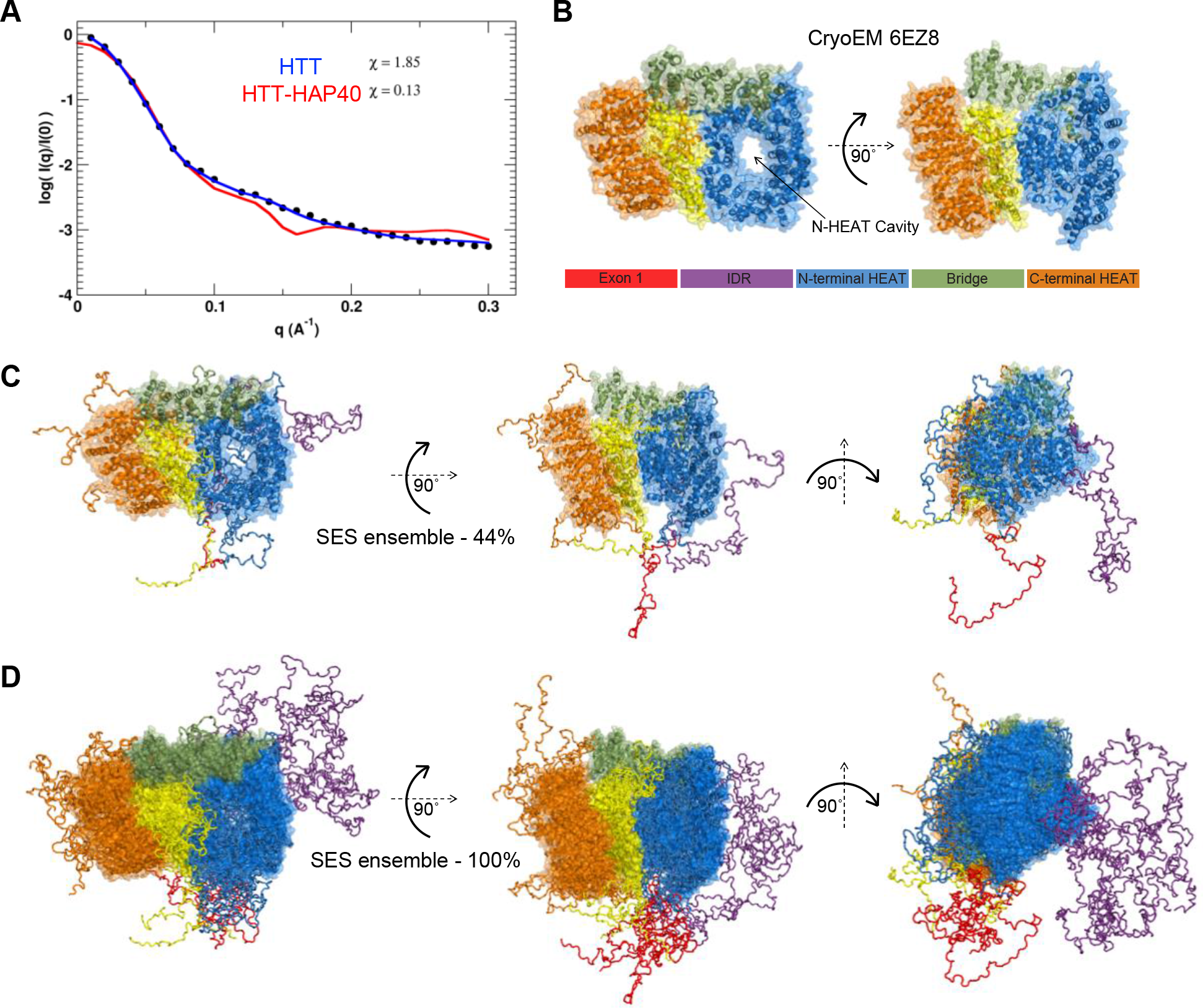
Fitting SAXS data to structural models of HTT-HAP40 complex. **A**) Experimental SAXS profiles (black circles) plotted with the theoretical profile for the EM structure of the HTT-HAP40 complex (red line) and the predicted profile averaged over the optimal Sparse Ensemble Selection (SES) ensemble (blue line). **B**) Surface representation of the EM structure as well as **C**) the most populated (~44 %) solution model and **D**) the complete ensemble of the HTT-HAP40 complex are shown with surface representation of the globular regions and backbone trace of the flexible regions (colour key as in panel B).

The most populated model (44%) in this ensemble (**Figure 8C**) shows extensive protruding regions of disorder extending out from the rigid complex core indicating that the overall envelope of the protein is likely to be larger than that calculated from the cryo-EM structure (**Figure 8B**). For many of the HEAT repeats, the disordered regions of connecting sequence are not see in the cryo-EM structure but the molecular modelling we have completed allows us to visualise how these might be arranged with respect to the rigid HEAT repeat core structure. The complete SES ensemble (**Figure 8D**) further indicates how, in particular, the IDR and exon1 region of huntingtin are probably very structurally heterogenous and dynamic in their conformation and are able to extend away from the more rigid core of the structure in many different arrangements due to their inherent flexibility. The extension of both exon1 and the IDR away from the core HTT-HAP40 complex is consistent with the accessibility of these domains to enzymes capable of post-translational modification.

A key feature of the cryo-EM structure is the large cavity which extends through the N-terminal HEAT repeat domain. This cavity is approximately the same diameter as a double stranded DNA helix and it is tempting to envision a potential nucleic acid binding role of this region of the HTT protein, especially given to the functional links between HTT and DNA damage repair (36, 37). At the current resolution of the structural information, it is also difficult to analyse potential surface charge or “greasy” surface residue hot spots which could indicate interaction sites. However, our SES ensemble model indicates how this cavity could be capped by certain conformational states of disordered loop regions, not resolved in the cryo-EM structure. These loops could act as gatekeepers to any binding partner, nucleic acid or protein, from accessing this cavity. Similarly, an apparent cavity in the side of the N-terminal HEAT repeat domain in the cryo-EM model, may also be capped by a flexible protein chain. Expansion of the polyQ region seems unlikely to affect the global structure of huntingtin given that exon1 is distal from the rigid HEAT repeat domains. Therefore, the mechanism by which the polyQ expansion affects huntingtin protein structure and function remains a question for future structural studies.

## DISCUSSION

We have generated a resource of 28 different HTT expression constructs which allow the generation of purified HTT samples of different polyQ lengths and affinity tags from three different expression systems. All constructs are available through Addgene, including the entry vector which will allow other researchers to make additional CAG expansion forms of the HTT gene should they require them. Our expression constructs permit facile scaling of culture volumes to enable the purification of milligram quantities of wild-type HTT protein from both insect and mammalian cells as well as substantial production of various polyQ expanded HTT species.

These HTT proteins from different expression systems are modified with PTMs previously described in the literature as well as additional modifications of unknown physiological relevance but that do not seem to alter HTT function in its ability to form a complex with HAP40. We describe HTT protein methylation for the first time, a modification conferred by various protein methyltransferases, many with links to neurodegeneration (38). Further characterisation of these modifications and their function could open up novel avenues to understanding HTT protein structure-function. The constructs cloned may also be used in future studies for co-expression with modifying enzymes to make highly site-specific PTMs on the sample as well as to test how certain enzymes might function on HTT.

Our work characterising the biophysical properties of HTT confirms that the protein is not monodisperse or homogenous when purified in its apo form. Coexpression and purification with HAP40 allows purification of a more monodisperse protein sample, rendering it amenable to more detailed structural analysis, as performed previously by cryo-EM (23). It is unclear whether HAP40 is a constitutive binder of HTT in physiological settings although its effects on the biophysical characteristics of the HTT protein are clearly significant. Interestingly, very few HTT protein-protein interaction studies have identified HAP40 as an interacting protein of HTT and it is only identified in the published literature in a small number of articles (23, 39) compared to the multiple extensive HTT interaction network publications (41, 42).

Due to the high yield of both HTT and HTT-HAP40 proteins from our toolkit of resources, were able to conduct preliminary biophysical analyses of these samples. Our SAXS analysis in tandem with molecular dynamics simulations permitted the generation of an SES ensemble, representing a possible solution structure of the HTT-HAP40 complex. This model gives insight into how HTT is post-translationally modified at flexible and accessible regions of sequence and suggests potential regulatory mechanisms such as steric capping of binding regions of the protein. Our SAXS model serves as an important resource for understanding the complete HTT-HAP40 complex, and should help prevent misinterpretation of certain features of the cryo-EM structure which lacks ~26% of the protein molecule. In particular, the exon1 region of HTT is distal to the complex core and polyQ expansion does not affect HTT thermal stability as shown by DSLS. Therefore, it is likely that the effect of polyQ expansion on HTT structure-function is more nuanced. Both exon1 and the IDR have many of the hallmarks of interacting domains observed for intrinsically disordered protein regions (43), as they are heavily modified by PTMs, are conformationally flexible and contain regions of charged amino acids (i.e. nearly 20 % of residues in the IDR are negatively charged). The purported role of huntingtin as a scaffold protein could be explained through dynamic protein-protein interactions mediated through these regions of the structure (40, 41).

The precise molecular function of unexpanded HTT remains elusive, so it is unclear how the polyQ expansion may alter the HTT protein sufficiently to give rise to the wide-ranging biochemical dysfunction observed in HD models and patients. These reagents and the accompanying methods and validation for the production of HTT protein will provide an enabling framework for future research requiring purified HTT and its complexes for a wide range of polyQ lengths.

## EXPERIMENTAL PROCEDURES

### Cloning of HTT and HAP40 expression constructs

HTT expression constructs were assembled in two steps into the mammalian/insect cell vector pBMDEL, an unencumbered vector created for open distribution of these reagents. First, entry vectors for N-terminal Flag tagged and C-terminal FLAG-tagged HTT (amino acids 1-3144) were constructed without the polyQ regions, amino acids 7-85. PCR products encoding wildtype HTT were amplified from cDNA (Kazusa Clone FHC15881) using primers N_int_FWD (ttaagaaggagatatactATGGACTACAAAGACGATGACGACAAGATGGCGACCCTGGAAAAGCgctGACCTTAGTCGCTAAcctgcaGGAGCCGCTGCACCGACCAAAG)/N_int_REV (gattggaagtagaggttttaGCAGGTGGTGACCTTGT GGAC) for the N-terminal FLAG-tagged HTT and C_int_FWD (ttaagaaggagatatactATGGCGACCCTGGAAAAGCgctGACCTTAGTCGCTAAcctgcaGGAGCCGCTGCACCGACCAAAG) / C_int_REV (gattggaagtagaggttttaCTTGTCGTCATCGTCTTTGTAGTCaccgcttccaccGCAGGTGGTGACCTTGTGGAC) for the C-terminal FLAG-tagged HTT. All PCR products were inserted using the In-Fusion cloning kit (Clontech) into the pBMDEL which had been linearized with BfuAI. Second, synthetic polyQ regions were inserted into the intermediate plasmids using the In-Fusion cloning kit. The polyQ regions were PCR amplified using the primers polyQP_Fwd (ATGGCGACCCTGGAAAAGCTG) / polyQP_Rev (TGGTCGGTGCAGCGGCTCCTC) from template plasmids CH00007 (Q23), CH00008 (Q73), and CH00065 (Q145) (all from Coriell Institute biorepository). PCR products were inserted into the intermediate vectors which had been linearized with Afe1 and Sbf1. Upon screening the assembled HTT expression constructs we found that our cloning method generated a range of polyQ lengths. We selected constructs with polyQ lengths from Q15 to Q145. The HTT coding sequences of intermediate and final expression constructs were confirmed by DNA sequencing. The sequences were confirmed by Addgene where these reagents have been deposited. HAP40 cDNA corresponding to amino acids 1-371 was subcloned into pFBOH-MHL expression vector using ligase independent cloning.

### HTT and HTT-HAP40 protein expression

The recombinant transfer vectors HTT pBMDEL and HAP40 pFBOH-MHL were transformed into DH10Bac E. coli cells (Invitrogen, Bac-to-Bac System) to generate recombinant Bacmid DNA. Sf9 cells (Invitrogen) were transfected with Bacmid DNA using jetPRIME® transfection reagent (PolyPlus Transfection, Cat. 89129-924), and recombinant baculovirus particles were recovered. The recombinant virus titre was sequentially amplified from P1 to P3 viral stocks for protein production in the Sf9 insect cells and EXPI293F mammalian cells.

Baculovirus-mediated expression of HTT in Insect Cells: Sf9 cells at a density of ~4.5 million cells per mL were infected with 8 mL of P3 recombinant baculovirus and grown at 130 rpm and 27 °C. HyQ SFX insect serum medium containing 10 μg/mL gentamicin was used as the culture medium. Infected cells were harvested when viability dropped to 80%–85%, normally after ~72 h post-infection. For HTT-HAP40 complex production, the same general protocol was followed but with a 3:1 ratio of HTT:HAP40 P3 recombinant baculovirus infection step.

Transduction of HTT in Mammalian Cells: EXPI293F cells (Thermo Fisher, Cat. A14527) were maintained in EXPI293 Expression Medium (Thermo Fisher, Cat. A1435102) in a humidified 8 % CO_2_ incubator at 37 °C and 125 rpm. Cells were transduced at a density of 2-3 million cells per mL culture. The transduction used recombinant baculoviruses of HTT constructs generated by transfecting Sf9 cells using Transfer vector pBMDEL and JetPRIME® transfection reagent (Cat. 89129-924). The volume of the virus added into the cells was at ratio as 6% of the total volume of the production. Infected cells were harvested after 7-10 days post-transduction depending on cell viability. Transient Transfection of HTT in Mammalian Cells: EXPI293F cells (Thermo Fisher, Cat. A14527) were maintained in EXPI293 Expression Medium (Thermo Fisher, Cat. A1435102) in a humidified 8 % CO_2_ incubator at 37 °C and 125 rpm. Cells were transfected at a density of 2-3 million cells per mL culture. FectoPRO® transfection reagent (VWR, Cat. 116-001) and plasmid pBMDEL-HTT DNA were separately diluted in serum-free OptiMEM complexation medium (Thermo Fisher, Cat. 31985062) at 10% of the total production volume in a ratio of 1 μg DNA to 1.2 μL FectoPro to 0.5 μL Booster per mL cell culture. After 5 min of incubation at room temperature, the transfection mixtures were combined and incubated for an additional 20 min. The FectoPRO® transfection reagent-DNA-OptiMEM mixture was then added to cells with an addition of FectoPRO® Booster in a ratio of 1 μg DNA to 1.2 μL FectoPro to 0.5 μL Booster per mL cell culture. The transfected cultures of EXPI293 cells were harvested after 72- 96 h post-transfection dependent cells density and viability.

### HTT and HTT-HAP40 protein purification

The same protocol was used to purify HTT from insect and mammalian cell culture, adapted from Huang *et al.* (2015) and Guo *et al.* (2018) (16, 23). Cell cultures were harvested by centrifugation at 4000 rpm, 20 mins, 4 °C (Beckman JLA 8.1000), washed in pre-chilled PBS and resuspended in 20 cell paste volumes of preparation buffer (50 mM Tris pH 8, 500 mM NaCl) and stored at −80 °C prior to purification. Cell suspensions were thawed and diluted to at least 50 times the cell paste volumes with prechilled buffer and supplemented with 1 mM PMSF, 1 mM benzamidine-HCl and 20 U/mL benzonase. NB: two freeze-thaw cycles of cell suspensions were found to be sufficient for cell lysis. The lysate was clarified by centrifugation at 14,000 rpm, 1 h, 4 °C (Beckman JLA 16.2500) and then bound to 0.1 cell paste volumes of anti-FLAG resin (Sigma M2) at 4 °C with rocking for 2 hours. Anti-FLAG resin was washed twice with the 100 cell paste volumes of buffer. HTT protein was eluted with 1 cell paste volume of buffer supplemented with 250 μg/mL 3xFLAG peptide (Chempep) run twice over the anti-FLAG resin. Residual HTT protein was washed from the beads with 0.5 cell paste volume of buffer. The sample was spin concentrated with MWCO 100,000. Depending on the protein yield, the sample was run as one or more sample runs on Superose 6 10/300 GL column in size-exclusion chromatography buffer (20 mM HEPES pH 7.5, 300 mM NaCl, 5 % (v/v) glycerol, 1 mM TCEP) at 0.4 mL/min ensuring no more than 2 mg of protein was applied per run to minimize protein aggregation. For HTT-HAP40, the same protocol was followed except for using preparation buffer with just 300 mM NaCl, and an additional step where FLAG elution was rocked with 2 mL equilibrated Ni-NTA resin for 30 mins before washing in preparation buffer and then elution with buffer supplemented with 300 mM imidazole prior to the size-exclusion chromatography step.

### SDS-PAGE and Western Blot analysis

SDS-PAGE and Western blot analysis were performed according to standard protocols. In brief, purified proteins were denatured in sample buffer (50 mM Tris HCl, 0.1 M DTT, 2% SDS, 0.1% bromophenol blue and 10% glycerol, pH 6.8) at 98 °C for 5 min, followed by sodium dodecyl sulfate polyacrylamide gel electrophoresis (SDS-PAGE). After electrophoresis, the gel was either directly stained with Coomassie Blue or subjected to Western blot analysis. For Western blot analysis, proteins were transferred onto PVDF membranes. The primary antibody used was anti-HTT (Abcam, ab109115, 1:5000) and the secondary antibody used was IRDye® 800CW anti-rabbit IgG (LI-COR, 926-32211, 1:5000). Membranes were visualised on an Odyssey® CLx Imaging System (LI-COR).

### HTT mass spectrometry

All data was acquired on an Agilent 1260 capillary HPLC system coupled to an Agilent Q-TOF 6545 mass spectrometer via the Dual Agilent Jetstream ion source.

Bottom-up proteomics for sequence coverage and PTM analysis: Proteins were processed according to established protocols (42). Briefly, proteins were reduced with DTT (10 mM final concentration) for 30 minutes at room temperature, alkylated with iodoacetamide (55 mM final concentration) for 30 minutes at room temperature, and incubated with trypsin (6 μL, 0.2 mg/mL) overnight at 37 °C. The digests acidified to pH 2 in hydrochloric acid and desalted on-column (by diverting the first two minutes to waste), before analysis. Peptides were separated on a C18 Advance BioPeptide column (2.1×150 mm 2.7 micron particles) at a flow rate of 400 microliters/min and an operating pressure of 4,700 psi. Peptides were eluted using a gradient from 100% solvent A (98:2 H_2_O:ACN with 1% formic acid) to 50% B (96:4 ACN:H2O with 1% formic acid) in 80 minutes. Mass spectra were acquired from m/z 300–1700 at a rate of 8 spectra per second. The tandem mass spectra were acquired in automated MS/MS mode from m/z 100-1500 with an acquisition rate of 3 spectra per second. The top ten precursors were selected and sorted by abundance only. Collision-induced dissociation was done using all ions at 4*(m/z)/100-1 and −5.

Data analysis: Raw data was processed using PEAKS Studio 8.5 (build 20171002) and the reference complete human proteome FASTA file (Uniprot). Cysteine carbamidomethylation was selected as a fixed modification, and methionine oxidation and N/Q deamidation as variable modifications. A minimum peptide length of five, a maximum of three missed cleavage sites, and a maximum of three labelled amino acids per peptide were employed.

### HTT sequence disorder prediction

Disorder prediction was performed using IUPred (33, 43). A threshold of 0.5 was used to define disordered or ordered regions, with predicted disordered regions shaded in light red. Further sequence analysis of HTT was performed using the sequence analysis tool local CIDER (32). Hydrophobicity was calculated using a normalized Kyte-Doolitle scale (43, 44). Resolved structure and domain annotations based on the solved cryo-EM structure of HTT in complex with HAP40 (23).

### Differential static light scattering (DSLS)

The thermal stability of HTT^1-3144^ Q19, Q23, Q42 and Q54 as well as HTT^1-3144^-HAP40^1-371^ Q23 and Q54 samples were analysed by DSLS using StarGazer. With four repeats per sample, HTT and HTT-HAP40 proteins at 1 mg/mL in 20 mM HEPES pH 7.5, 300 mM NaCl, 5 % (v/v) glycerol, 1 mM TCEP were heated from 20 °C to 85 °C. Protein aggregation was monitored using a CCD camera. The temperature of aggregation (T_agg_) was analysed and fitted as described previously (35).

### Size Exclusion Chromatography Multi-Angle Light Scattering (SEC-MALS)

The absolute molar masses and mass distributions of purified protein samples of HTT^1-3144^ Q23 and HTT^1-3144^ Q23-HAP40^1-371^ complex with C-terminal FLAG tag at 1 mg/mL were determined using SEC-MALS. Samples were injected through a Superose 6 10/300 GL column (GE Healthcare Life Sciences) equilibrated in 20 mM HEPES pH 7.5, 300 mM NaCl, 5 % (v/v) glycerol, 1 mM TCEP followed in-line by a Dawn Heleos-II light scattering detector (Wyatt Technologies) and a 2414 refractive index detector (Waters). Molecular mass calculations were performed using ASTRA 6.1.1.17 (Wyatt Technologies) assuming a dn/dc value of 0.185 ml/g.

### SAXS data collection and analysis

SAXS measurements were carried out at the beamline 12-ID-B of the Advanced Photon Source, Argonne National Laboratory. The energy of the X-ray beam was 14 Kev (wavelength λ=0. 8856 Å), and two setups (small- and wide-angle X-ray scattering, SAXS and WAXS) were used simultaneously to cover scattering *q* ranges of 0.006 < *q* < 2.6 Å^−1^, where q = (4π/λ)sinθ, and 2θ is the scattering angle. Thirty two-dimensional images were recorded for each buffer or sample solutions using a flow cell, with the accumulated exposure time of 0.8-2 seconds to reduce radiation damage and obtain good statistics. No radiation damage was observed as confirmed by the absence of systematic signal changes in sequentially collected X-ray scattering images. The 2D images were corrected and reduced to 1D scattering profiles using the Matlab software package at the beamlines. The 1D SAXS profiles were grouped by sample and averaged.

Concentration-series measurements for each sample were carried out at 300 K. We used HTT concentrations of 1.0, 2.0, and 4.0 mg/mL, and complex HTT/HAP40 concentrations of 0.5, 1.0, and 2.0 mg/mL, respectively in 20 mM HEPES pH 7.5, 300 mM NaCl, 5 % (v/v) glycerol, 1 mM TCEP. The data were processed and analyzed with ATSAS program package (45). The scattering profile of the protein was calculated by subtracting the background buffer contribution from the sample-buffer profile and the difference data were extrapolated to zero solute concentration by standard procedures. Guinier analysis and the experimental radius of gyration (*R*_g_) estimation from the data of infinite dilution were performed using PRIMUS. The pair distance distribution function (PDDF), p(r), and the maximum dimension of the protein, *D*_max_, in real space was calculated with the indirect Fourier transform using program GNOM (46). The molecular weights were estimated separately based on volume calculated by SAXMoW (47), and Volume of correlation Vc calculated by DATVC in q range of 0 < q < 0.3 Å^−1^. The theoretical scattering intensity of a structural model was calculated and fitted to the experimental scattering intensity using CRYSOL and FoXS (50) programs.

### Fitting structural ensemble to SAXS data

The SAXS data indicate that the HTT/HAP40 complex possess some degree of flexibility. The known EM structure of the complex (PDB: 6ez8) is missing ~26% of the residues and it does not fit the SAXS data. We assume that the residues with known coordinates form a quasi-rigid part of the complex while the residues with missing coordinates are flexible. We performed coarse-grained MD simulations to generate the initial ensemble of possible conformations of the complex. MD trajectory of 1000ns was generated at 300K and theoretical scattering profiles in the *q* range 0 < *q* < 0.3 Å^−1^ for 5000 frames taken from the trajectory were calculated using FoXS. The calculated scattering curves were averaged over the entire ensemble of structures using the optimal weights for each ensemble member obtained with SES method (35), and this average profile was compared with the experimental scattering data.

### Coarse-grained molecular dynamics simulations

We used a coarse-grained model of HTT/HAP40 protein complex in order to enhance the sampling efficiency in the conformational space of the complex. In this model, amino acid residues in the proteins are represented as single beads located at their C_α_ positions and interacting via appropriate bonding, bending, torsion-angle, and non-bonding potential. A Gō-like model of Clementi and Onuchic (51) was employed to maintain the structured, globular domains as quasi-rigid in the simulation. For flexible regions, we adopt simple model in which adjacent amino acids beads are joined together into a polymer chain by means of virtual bond and angle interactions with a quadratic potential.

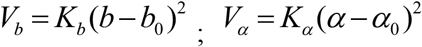

with the constants *Kb* = 50 *kcal/mol* and *K*_*α*_ = 1.75 *kcal/mol* and the equilibrium values *b*_0_ = 3.8 Å and *α*_0_ = 112° for bonds and angles, respectively. The excluded volume between nonbonded beads was treated with pure repulsive potential

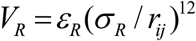

where *r*_*ij*_ is the inter-bead distance, *σ*_*R*_ = 4 Å, and *ε*_*R*_ = 2.0 *kcal/mol*.

The interaction between quasi-rigid domains is modeled with the residue-specific pair interaction potentials that combine short-range interactions with the long-range electrostatics as it described (52, 53). The short-range interaction is given by a Lennard-Jones 12-10-6 -type potential and simple Debye-Hückel-type potential is used for the electrostatics interactions (53). In this study we used the dielectric constant of 80 and the Debye screening length of 10 Å, which corresponds to a salt concentration of about 100 mM. In-house software was developed and used to carry out constant temperature molecular dynamics simulations of the coarse-grained model described above.

## ACKNOWLEDGEMENTS

We thank Dr. Xiaobing Zuo (Argonne National Laboratory) for expert support with SAXS measurements, and acknowledge the use of the SAXS Core facility of Center for Cancer Research (CCR), National Cancer Institute (NCI) which is funded by Frederick National Laboratory for Cancer Research under contract HHSN261200800001E and the intramural research program of the NIH, NCI, CCR. The content of this publication does not necessarily reflect the views or policies of the Department of Health and Human Services, nor does mention of trade names, commercial products or organizations imply endorsement by the US Government. This research used 12-ID-B beamline of the Advanced Photon Source, a U.S. Department of Energy (DOE) Office of Science User Facility operated for the DOE Office of Science by Argonne National Laboratory under Contract No. DE-AC02-06CH11357. We alsoWe acknowledge Dr. Pravin Mahajan who constructed the pBMDEL vector. This research was supported by a Huntington’s Disease Society of America Berman Topper Career Development Fellowship (RH), the Natural Sciences and Engineering Research Council of Canada (CHA) (grant RGPIN-2015-05939), Huntington Society of Canada (CHA), CHDI Foundation (LTS, CHA) and the SGC, a registered charity (number 1097737) that receives funds from AbbVie, Bayer Pharma AG, Boehringer Ingelheim, Canada Foundation for Innovation, Eshelman Institute for Innovation, Genome Canada through Ontario Genomics Institute [OGI-055], Innovative Medicines Initiative (EU/EFPIA) [ULTRA-DD grant no. 115766], Janssen, Merck KGaA, Darmstadt, Germany, MSD, Novartis Pharma AG, Ontario Ministry of Research, Innovation and Science (MRIS), Pfizer, São Paulo Research Foundation-FAPESP, Takeda, and Wellcome. ASH is a postdoctoral fellow in the laboratory of R.V. Pappu at Washington University in St. Louis and is funded by the Human Frontiers Science Program (grant RGP0034/2017).

## DATA AND ANALYSIS FILES

All contributing data and analysis files for this manuscript can be found under Creative Commons Attribution 4.0 International license in Zenodo: https://zenodo.org/record/2169035. Mass spectrometry data is also available through PRIDE - accession PXD010865.

