## Supplementary Information for "Design and characterisation of mutant and wild-type huntingtin proteins produced from a toolkit of scalable eukaryotic expression systems"

**Running title:** A toolkit of HTT protein resources

† Present address: Custom Biologics, 115 Skyway Ave, Toronto, Ontario M9W 4Z4, Canada
‡ Present address: The Tau Consortium, The Rainwater Charitable Foundation, 777 Main St, Ste 2250, Fort Worth, Texas 76102, USA

*To whom correspondence should be addressed:

Rachel J. Harding,


Cheryl H. Arrowsmith


**Table of contents:**

p S-2 - Table S1. Database search results for HTT$^{1-3144}$ Q23 expressed in EXPI293F cells.
p S-3 - Figure S1. Western blot analysis of HTT samples from different expression systems.
p S-4 - Figure S2. Mass spectrometry sequence coverage maps of Sf9 HTT$^{1-3144}$ Q23
p S-5 - Figure S3. Mass spectrometry sequence coverage maps of EXPI293F HTT$^{1-3144}$ Q23
p S-6 - Figure S4. Exemplary spectra of peptides identified after digesting HTT$^{1-3144}$ Q23 from Sf9 with pepsin.
p S-8 - Figure S5. Exemplary spectra for peptides identified after digesting HTT$^{1-3144}$ Q23 from Sf9 with trypsin or lysargiNase.
p S-13 - Figure S6. Exemplary spectra of peptides identified after digesting HTT$^{1-3144}$ Q23 from EXPI293F with trypsin.
p S-19 - Figure S7. Mapping HTT posttranslational modifications identified from HTT$^{1-3144}$ samples from Sf9 and EXPI293F cells onto the HTT structure.
p S-20 – References

**Table S1. Database search results for HTT$^{1-3144}$ Q23 expressed in EXPI293F cells.** Proteins with five or more total no. spectra are listed. Of the peptides which do not correspond to HTT protein, most either have very high scores in the CRAPome (1), suggesting that these are non-specific contaminants of the co-immunoprecipitation step, not true HTT interactors, or they are very low abundance.

| Protein Accession | Total no. spectra | No. unique peptides | Protein Accession | Total no. spectra | No. unique peptides | Protein Accession | Total no. spectra | No. unique peptides |
|---|---|---|---|---|---|---|---|---|
| P42858\|HD | 7681 | 1150 | P0C0S5\|H2AZ | 14 | 4 | O60391\|NMD3B | 8 | 2 |
| Q13748\|TBA3C | 175 | 29 | P0C0S8\|H2A1 | 14 | 4 | O75534\|CSDE1 | 8 | 2 |
| Q13885\|TBB2A | 111 | 26 | P16104\|H2AX | 14 | 4 | P49137\|MAPK2 | 8 | 2 |
| Q9BVA1\|TBB2B | 111 | 26 | P20671\|H2A1D | 14 | 4 | P62269\|RS18 | 8 | 2 |
| P07900\|HS90A | 110 | 24 | P34931\|HS71L | 14 | 5 | Q16695\|H31T | 8 | 2 |
| Q6PEY2\|TBA3E | 64 | 13 | Q16777\|H2A2C | 14 | 4 | Q8NCG5\|CHST4 | 8 | 2 |
| Q9BYE2\|TMPSD | 56 | 7 | Q6FI13\|H2A2A | 14 | 4 | Q8WUA7\|TB22A | 8 | 2 |
| Q9NY65\|TBA8 | 54 | 9 | Q71UI9\|H2AV | 14 | 4 | Q9NRC6\|SPTN5 | 8 | 2 |
| Q71U36\|TBA1A | 51 | 12 | Q7L7L0\|H2A3 | 14 | 4 | Q9NRD9\|DUOX1 | 8 | 2 |
| P04350\|TBB4A | 48 | 12 | Q93077\|H2A1C | 14 | 4 | O15381\|NVL | 7 | 2 |
| P60709\|ACTB | 45 | 13 | Q96KK5\|H2A1H | 14 | 4 | Q15021\|CND1 | 7 | 2 |
| P11142\|HSP7C | 44 | 12 | Q96QV6\|H2A1A | 14 | 4 | Q5T9S5\|CCD18 | 7 | 2 |
| P63261\|ACTG | 44 | 13 | Q99878\|H2A1J | 14 | 4 | Q5VZP5\|DUS27 | 7 | 2 |
| P23396\|RS3 | 43 | 11 | Q9BTM1\|H2AJ | 14 | 4 | Q6WRI0\|IGS10 | 7 | 2 |
| P54652\|HSP72 | 43 | 12 | Q9H2M9\|RBGPR | 14 | 2 | Q709C8\|VP13C | 7 | 2 |
| P15880\|RS2 | 38 | 11 | Q9NY93\|DDX56 | 14 | 3 | Q96JC1\|VPS39 | 7 | 2 |
| Q13509\|TBB3 | 36 | 8 | Q9P225\|DYH2 | 14 | 4 | Q9BSJ2\|GCP2 | 7 | 2 |
| tr\|A0A0B4J269 | 32 | 7 | Q9UKN7\|MYO15 | 14 | 3 | Q9P1Y6\|PHRF1 | 7 | 2 |
| P0DMV8\|HS71A | 31 | 9 | Q58FF7\|H90B3 | 13 | 5 | Q9P1Z3\|HCN3 | 7 | 2 |
| P0DMV9\|HS71B | 31 | 9 | Q8IVL1\|NAV2 | 13 | 3 | Q9P2E5\|CHPF2 | 7 | 2 |
| P07437\|TBB5 | 30 | 7 | Q9H853\|TBA4B | 13 | 3 | Q9UL51\|HCN2 | 7 | 2 |
| Q9BWH6\|RPAP1 | 27 | 4 | P02549\|SPTA1 | 12 | 2 | Q9Y3Q4\|HCN4 | 7 | 2 |
| Q5VYK3\|ECM29 | 26 | 6 | P62805\|H4 | 12 | 2 | A6NE52\|WDR97 | 6 | 1 |
| Q8WXG9\|GPR98 | 26 | 4 | Q14568\|HS902 | 12 | 5 | O43314\|VIP2 | 6 | 1 |
| Q562R1\|ACTBL | 24 | 7 | Q5CZC0\|FSIP2 | 12 | 5 | O43896\|KIF1C | 6 | 2 |
| Q9UPN3\|MACF1 | 23 | 4 | Q5S007\|LRRK2 | 12 | 2 | P05164\|PERM | 6 | 2 |
| A6NKT7\|RGPD3 | 22 | 6 | Q7Z5J8\|ANKAR | 12 | 3 | P08670\|VIME | 6 | 1 |
| P17066\|HSP76 | 21 | 7 | Q8TE73\|DYH5 | 12 | 6 | P19823\|ITIH2 | 6 | 2 |
| Q03001\|DYST | 21 | 3 | Q96JB1\|DYH8 | 12 | 4 | Q13608\|PEX6 | 6 | 1 |
| P08238\|HS90B | 20 | 7 | Q96PX9\|PKH4B | 12 | 2 | Q13683\|ITA7 | 6 | 1 |
| Q9BQG0\|MBB1A | 20 | 5 | Q9UDT6\|CLIP2 | 12 | 3 | Q2LD37\|K1109 | 6 | 1 |
| P11021\|GRP78 | 19 | 6 | O60890\|OPHN1 | 11 | 3 | Q5JV73\|FRPD3 | 6 | 3 |
| Q58FG1\|HS904 | 19 | 4 | Q02224\|CENPE | 11 | 3 | Q5JWR5\|DOP1 | 6 | 1 |
| Q8NET8\|TRPV3 | 19 | 3 | Q6ZQQ6\|WDR87 | 11 | 2 | Q6N069\|NAA16 | 6 | 2 |
| O60814\|H2B1K | 18 | 3 | Q7Z333\|SETX | 11 | 4 | Q86XA9\|HTR5A | 6 | 1 |
| P06899\|H2B1J | 18 | 3 | Q86UQ4\|ABCAD | 11 | 4 | Q8NCM8\|DYHC2 | 6 | 1 |
| P23527\|H2B1O | 18 | 3 | Q9Y4F4\|TGRM1 | 11 | 3 | Q8TD57\|DYH3 | 6 | 2 |
| P33778\|H2B1B | 18 | 3 | P21817\|RYR1 | 10 | 2 | Q92997\|DVL3 | 6 | 1 |
| P57053\|H2BFS | 18 | 3 | Q12955\|ANK3 | 10 | 5 | Q96A08\|H2B1A | 6 | 1 |
| P58876\|H2B1D | 18 | 3 | Q1XH10\|SKDA1 | 10 | 2 | Q96QT4\|TRPM7 | 6 | 1 |
| P62807\|H2B1C | 18 | 3 | Q5T1B0\|AXDN1 | 10 | 2 | Q99996\|AKAP9 | 6 | 3 |
| Q16778\|H2B2E | 18 | 3 | Q86U86\|PB1 | 10 | 3 | Q9HCK8\|CHD8 | 6 | 3 |
| Q5QNW6\|H2B2F | 18 | 3 | Q86VI3\|IQGA3 | 10 | 3 | Q9NYC9\|DYH9 | 6 | 1 |
| Q8N257\|H2B3B | 18 | 3 | Q8IZD9\|DOCK3 | 10 | 3 | Q9P2D1\|CHD7 | 6 | 3 |
| Q93079\|H2B1H | 18 | 3 | Q13136\|LIPA1 | 9 | 4 | P09131\|P3 | 5 | 1 |
| Q99877\|H2B1N | 18 | 3 | Q15751\|HERC1 | 9 | 3 | Q13576\|IQGA2 | 5 | 2 |
| Q99879\|H2B1M | 18 | 3 | Q6PI48\|SYDM | 9 | 3 | Q15746\|MYLK | 5 | 1 |
| Q99880\|H2B1L | 18 | 3 | Q9HBJ7\|UBP29 | 9 | 3 | Q86VV8\|RTTN | 5 | 3 |
| Q12931\|TRAP1 | 16 | 5 | Q9P0L2\|MARK1 | 9 | 1 | Q8NBX0\|SCPDL | 5 | 2 |
| Q8IVF2\|AHNK2 | 16 | 1 | Q9P217\|ZSWM5 | 9 | 3 | Q8TD26\|CHD6 | 5 | 2 |
| O14647\|CHD2 | 15 | 2 | tr\|Q5TEC6\|Q5TEC6 | 9 | 3 | Q9ULT8\|HECD1 | 5 | 3 |
| P04908\|H2A1B | 14 | 4 | O15078\|CE290 | 8 | 3 | | | |

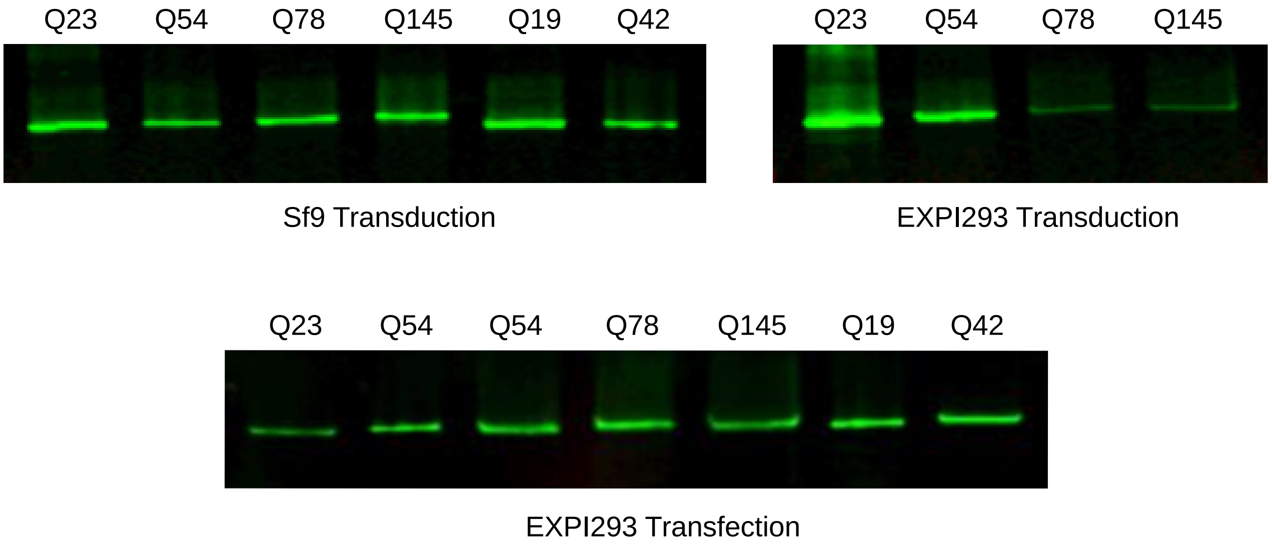


**Figure S1. Western blot analysis of HTT samples from different expression systems.** C-terminally FLAG-tagged HTT derived from EXPI293F expression, either by transient transfection or baculoviral transduction in EXPI293F cells or baculoviral transduction in Sf9 cells, were subject to Western Blot analysis with ab109115 which binds an epitope at amino acids 1-100.

A)

```
   1 MATLEKLMKA FESLKSFQQQ QQQQQQQQQQ QQQQQQQQPP PPPPPPPPPQ LPQPPPQAQP LLPQPQPPPP PPPPPPGPAV AEEPLHRPKK ELSATKKDRV
 101 NHCLTICENI VAQSVRNSPE FQKLLGIAME LFLLCSDDAE SDVRMVADEC LNKVIKALMD SNLPRLQLEL YKEIKKNGAP RSLRAALWRF AELAHLVRPQ
 201 KCRPYLVNLL PCLTRTSKRP EESVQETLAA AVPKIMASFG NFANDNEIKV LLKAFIANLK SSSPTIRRTA AGSAVSICQH SRRTQYFYSW LLNVLLGLLV
 301 PVEDEHSTLL ILGVLLTLRY LVPLLQQQVK DTSLKGSFGV TRKEMEVSPS AEQLVQVYEL TLHHTQHQDH NVVTGALELL QQLFRTPPPE LLQTLTAVGG
 401 IGQLTAAKEE SGGRSRSGSI VELIAGGGSS CSPVLSRKQK GKVLLGEEEA LEDDSESRSD VSSSALTASV KDEISGELAA SSGVSTPGSA GHDIITEQPR
 501 SQHTLQADSV DLASCDLTSS ATDGDEEDIL SHSSSQVSAV PSDPAMDLND GTQASSPISD SSQTTTEGPD SAVTPSDSSE IVLDGTDNQY LGLQIGQPQD
 601 EDEEATGILP DEASEAFRNS SMALQQAHLL KNMSHCRQPS DSSVDKFVLR DEATEPGDQE NKPCRIKGDI GQSTDDDSAP LVHCVRLLSA SFLLTGGKNV
 701 LVPDRDVRVS VKALALSCVG AAVALHPESF FSKLYKVPLD TTEYPEEQYV SDILNYIDHG DPQVRGATAI LCGTLICSIL SRSRFHVGDW MGTIRTLTGN
 801 TFSLADCIPL LRKTLKDESS VTCKLACTAV RNCVMSLCSS SYSELGLQLI IDVLTLRNSS YWLVRTELLE TLAEIDFRLV SFLEAKAENL HRGAHHYTGL
 901 LKLQERVLNN VVIHLLGDED PRVRHVAAAS LIRLVPKLFY KCDQGQADPV VAVARDQSSV YLKLLMHETQ PPSHFSVSTI TRIYRGYNLL PSITDVTMEN
1001 NLSRVIAAVS HELITSTTRA LTFGCCEALC LLSTAFPVCI WSLGWHCGVP PLSASDESRK SCTVGMATMI LTLLSSAWFP LDLSAHQDAL ILAGNLLAAS
1101 APKSLRSSWA SEEEANPAAT KQEEVWPALG DRALVPMVEQ LFSHLLKVIN ICAHVLDDVA PGPAIKAALP SLTNPPSLSP IRRKGKEKEP GEQASVPLSP
1201 KKGSEASAAS RQSDTSGPVT TSKSSSLGSF YHLPSYLKLH DVLKATHANY KVTLDLQNST EKFGGFLRSA LDVLSQILEL ATLQDIGKCV EEILGYLKSC
1301 FSREPMMATV CVQQLLKTLF GTNLASQFDG LSSNPSKSQG RAQRLGSSSV RPGLYHYCFM APYTHFTQAL ADASLRNMVQ AEQENDTSGW FDVLQKVSTQ
1401 LKTNLTSVTK NRADKNAIHN HIRLFEPLVI KALKQYTTTT CVQLQKQVLD LLAQLVQLRV NYCLLDSDQV FIGFVLKQFE YIEVGQFRES EAIIPNIFFF
1501 LVLLSYERYH SKQIIGIPKI IQLCDGIMAS GRKAVTHAIP ALQPIVHDLF VLRGTNKADA GKELETQKEV VVSMLLRLIQ YHQVLEMFIL VLQQCHKENE
1601 DKWKRLSRQI ADIILPMLAK QQMHIDSHEA LGVLNTLFEI LAPSSLRPVD MLLRSMFVTP NTMASVSTVQ LWISGILAIL RVLISQSTED IVLSRIQELS
1701 FSPYLISCTV INRLRDGDST STLEEHSEGK QIKNLPEETF SRFLLQLVGI LLEDIVTKQL KVEMSEQQHT FYCQELGTLL MCLIHIFKSG MFRRITAAAT
1801 RLFRSDGCGG SFYTLDSLNL RARSMITTHP ALVLLWCQIL LLVNHTDYRW WAEVQQTPKR HSLSSTKLLS PQMSGEEEDS DLAAKLGMCN REIVRRGALI
1901 LFCDYVCQNL HDSEHLTWLI VNHIQDLISL SHEPPVQDFI SAVHRNSAAS GLFIQAIQSR CENLSTPTML KKTLQCLEGI HLSQSGAVLT LYVDRLLCTP
2001 FRVLARMVDI LACRRVEMLL AANLQSSMAQ LPMEELNRIQ EYLQSSGLAQ RHQRLYSLLD RFRLSTMQDS LSPSPPVSSH PLDGDGHVSL ETVSPDKDWY
2101 VHLVKSQCWT RSDSALLEGA ELVNRIPAED MNAFMMNSEF NLSLLAPCLS LGMSEISGGQ KSALFEAARE VTLARVSGTV QQLPAVHHVF QPELPAEPAA
2201 YWSKLNDLFG DAALYQSLPT LARALAQYLV VVSKLPSHLH LPPEKEKDIV KFVVATLEAL SWHLIHEQIP LSLDLQAGLD CCCLALQLPG LWSVVSSTEF
2301 VTHACSLIYC VHFILEAVAV QPGEQLLSPE RRTNTPKAIS EEEEEVDPNT QNPKYITAAC EMVAEMVESL QSVLALGHKR NSGVPAFLTP LLRNIIISLA
2401 RLPLVNSYTR VPPLVWKLGW SPKPGGDFGT AFPEIPVEFL QEKEVFKEFI YRINTLGWTS RTQFEETWAT LLGVLVTQPL VMEQEESPPE EDTERTQINV
2501 LAVQAITSLV LSAMTVPVAG NPAVSCLEQQ PRNKPLKALD TRFGRKLSII RGIVEQEIQA MVSKRENIAT HHLYQAWDPV PSLSPATTGA LISHEKLLLQ
2601 INPERELGSM SYKLGQVSIH SVWLGNSITP LREEEWDEEE EEEADAPAPS SPPTSPVNSR KHRAGVDIHS CSQFLLELYS RWILPSSSAR RTPAILISEV
2701 VRSLLVVSDL FTERNQFELM YVTLTELRRV HPSEDEILAQ YLVPATCKAA AVLGMDKAVA EPVSRLLEST LRSSHLPSRV GALHGVLYVL ECDLLDDTAK
2801 QLIPVISDYL LSNLKGIAHC VNIHSQQHVL VMCATAFYLI ENYPLDVGPE FSASIIQMCG VMLSGSEEST PSIIYHCALR GLERLLLSEQ LSRLDAESLV
2901 KLSVDRVNVH SPHRAMAALG LMLTCMYTGK EKVSPGRTSD PNPAAPDSES VIVAMERVSV LFDRIRKGFP CEARVVARIL PQFLDDFFPP QDIMNKVIGE
3001 FLSNQQPYPQ FMATVVYKVF QTLHSTGQSS MVRDWVMLSL SNFTQRAPVA MATWSLSCFF VSASTSPWVA AILPHVISRM GKLEQVDVNL FCLVATDFYR
3101 HQIEEELDRR AFQSVLEVVA APGSPYHRLL TCLRNVHKVT TC
```

B)

```
   1 MATLEKLMKA FESLKSFQQQ QQQQQQQQQQ QQQQQQQQPP PPPPPPPPPQ LPQPPPQAQP LLPQPQPPPP PPPPPPGPAV AEEPLHRPKK ELSATKKDRV
 101 NHCLTICENI VAQSVRNSPE FQKLLGIAME LFLLCSDDAE SDVRMVADEC LNKVIKALMD SNLPRLQLEL YKEIKKNGAP RSLRAALWRF AELAHLVRPQ
 201 KCRPYLVNLL PCLTRTSKRP EESVQETLAA AVPKIMASFG NFANDNEIKV LLKAFIANLK SSSPTIRRTA AGSAVSICQH SRRTQYFYSW LLNVLLGLLV
 301 PVEDEHSTLL ILGVLLTLRY LVPLLQQQVK DTSLKGSFGV TRKEMEVSPS AEQLVQVYEL TLHHTQHQDH NVVTGALELL QQLFRTPPPE LLQTLTAVGG
 401 IGQLTAAKEE SGGRSRSGSI VELIAGGGSS CSPVLSRKQK GKVLLGEEEA LEDDSESRSD VSSSALTASV KDEISGELAA SSGVSTPGSA GHDIITEQPR
 501 SQHTLQADSV DLASCDLTSS ATDGDEEDIL SHSSSQVSAV PSDPAMDLND GTQASSPISD SSQTTTEGPD SAVTPSDSSE IVLDGTDNQY LGLQIGQPQD
 601 EDEEATGILP DEASEAFRNS SMALQQAHLL KNMSHCRQPS DSSVDKFVLR DEATEPGDQE NKPCRIKGDI GQSTDDDSAP LVHCVRLLSA SFLLTGGKNV
 701 LVPDRDVRVS VKALALSCVG AAVALHPESF FSKLYKVPLD TTEYPEEQYV SDILNYIDHG DPQVRGATAI LCGTLICSIL SRSRFHVGDW MGTIRTLTGN
 801 TFSLADCIPL LRKTLKDESS VTCKLACTAV RNCVMSLCSS SYSELGLQLI IDVLTLRNSS YWLVRTELLE TLAEIDFRLV SFLEAKAENL HRGAHHYTGL
 901 LKLQERVLNN VVIHLLGDED PRVRHVAAAS LIRLVPKLFY KCDQGQADPV VAVARDQSSV YLKLLMHETQ PPSHFSVSTI TRIYRGYNLL PSITDVTMEN
1001 NLSRVIAAVS HELITSTTRA LTFGCCEALC LLSTAFPVCI WSLGWHCGVP PLSASDESRK SCTVGMATMI LTLLSSAWFP LDLSAHQDAL ILAGNLLAAS
1101 APKSLRSSWA SEEEANPAAT KQEEVWPALG DRALVPMVEQ LFSHLLKVIN ICAHVLDDVA PGPAIKAALP SLTNPPSLSP IRRKGKEKEP GEQASVPLSP
1201 KKGSEASAAS RQSDTSGPVT TSKSSSLGSF YHLPSYLKLH DVLKATHANY KVTLDLQNST EKFGGFLRSA LDVLSQILEL ATLQDIGKCV EEILGYLKSC
1301 FSREPMMATV CVQQLLKTLF GTNLASQFDG LSSNPSKSQG RAQRLGSSSV RPGLYHYCFM APYTHFTQAL ADASLRNMVQ AEQENDTSGW FDVLQKVSTQ
1401 LKTNLTSVTK NRADKNAIHN HIRLFEPLVI KALKQYTTTT CVQLQKQVLD LLAQLVQLRV NYCLLDSDQV FIGFVLKQFE YIEVGQFRES EAIIPNIFFF
1501 LVLLSYERYH SKQIIGIPKI IQLCDGIMAS GRKAVTHAIP ALQPIVHDLF VLRGTNKADA GKELETQKEV VVSMLLRLIQ YHQVLEMFIL VLQQCHKENE
1601 DKWKRLSRQI ADIILPMLAK QQMHIDSHEA LGVLNTLFEI LAPSSLRPVD MLLRSMFVTP NTMASVSTVQ LWISGILAIL RVLISQSTED IVLSRIQELS
1701 FSPYLISCTV INRLRDGDST STLEEHSEGK QIKNLPEETF SRFLLQLVGI LLEDIVTKQL KVEMSEQQHT FYCQELGTLL MCLIHIFKSG MFRRITAAAT
1801 RLFRSDGCGG SFYTLDSLNL RARSMITTHP ALVLLWCQIL LLVNHTDYRW WAEVQQTPKR HSLSSTKLLS PQMSGEEEDS DLAAKLGMCN REIVRRGALI
1901 LFCDYVCQNL HDSEHLTWLI VNHIQDLISL SHEPPVQDFI SAVHRNSAAS GLFIQAIQSR CENLSTPTML KKTLQCLEGI HLSQSGAVLT LYVDRLLCTP
2001 FRVLARMVDI LACRRVEMLL AANLQSSMAQ LPMEELNRIQ EYLQSSGLAQ RHQRLYSLLD RFRLSTMQDS LSPSPPVSSH PLDGDGHVSL ETVSPDKDWY
2101 VHLVKSQCWT RSDSALLEGA ELVNRIPAED MNAFMMNSEF NLSLLAPCLS LGMSEISGGQ KSALFEAARE VTLARVSGTV QQLPAVHHVF QPELPAEPAA
2201 YWSKLNDLFG DAALYQSLPT LARALAQYLV VVSKLPSHLH LPPEKEKDIV KFVVATLEAL SWHLIHEQIP LSLDLQAGLD CCCLALQLPG LWSVVSSTEF
2301 VTHACSLIYC VHFILEAVAV QPGEQLLSPE RRTNTPKAIS EEEEEVDPNT QNPKYITAAC EMVAEMVESL QSVLALGHKR NSGVPAFLTP LLRNIIISLA
2401 RLPLVNSYTR VPPLVWKLGW SPKPGGDFGT AFPEIPVEFL QEKEVFKEFI YRINTLGWTS RTQFEETWAT LLGVLVTQPL VMEQEESPPE EDTERTQINV
2501 LAVQAITSLV LSAMTVPVAG NPAVSCLEQQ PRNKPLKALD TRFGRKLSII RGIVEQEIQA MVSKRENIAT HHLYQAWDPV PSLSPATTGA LISHEKLLLQ
2601 INPERELGSM SYKLGQVSIH SVWLGNSITP LREEEWDEEE EEEADAPAPS SPPTSPVNSR KHRAGVDIHS CSQFLLELYS RWILPSSSAR RTPAILISEV
2701 VRSLLVVSDL FTERNQFELM YVTLTELRRV HPSEDEILAQ YLVPATCKAA AVLGMDKAVA EPVSRLLEST LRSSHLPSRV GALHGVLYVL ECDLLDDTAK
2801 QLIPVISDYL LSNLKGIAHC VNIHSQQHVL VMCATAFYLI ENYPLDVGPE FSASIIQMCG VMLSGSEEST PSIIYHCALR GLERLLLSEQ LSRLDAESLV
2901 KLSVDRVNVH SPHRAMAALG LMLTCMYTGK EKVSPGRTSD PNPAAPDSES VIVAMERVSV LFDRIRKGFP CEARVVARIL PQFLDDFFPP QDIMNKVIGE
3001 FLSNQQPYPQ FMATVVYKVF QTLHSTGQSS MVRDWVMLSL SNFTQRAPVA MATWSLSCFF VSASTSPWVA AILPHVISRM GKLEQVDVNL FCLVATDFYR
3101 HQIEEELDRR AFQSVLEVVA APGSPYHRLL TCLRNVHKVT TC
```

**Figure S2. Primary sequence coverage maps of Sf9 HTT$^{1\text{-}3144}$ Q23.** A) 97% coverage obtained by combining the database search results obtained from 5 enzymes (pepsin, WaLP, MaLP, lysargiNase, and trypsin), and B) 66% coverage obtained from 2 enzymes (lysargiNase and trypsin). NB: table displays HTT Q21 sequence as UniProt database used in sequence search.

A)

```
1    MATLEKLMKA FESLKSFQQQ QQQQQQQQQQ QQQQQQQQPP PPPPPPPPPQ LPQPPPQAQP LLPQPQPPPP PPPPPPGPAV AEEPLHRPKK ELSATKKDRV
101  NHCLTICENI VAQSVRNSPE FQKLLGIAME LFLLCSDDAE SDVRMVADEC LNKVIKALMD SNLPRLQLEL YKEIKKNGAP RSLRAALWRF AELAHLVRPQ
201  KCRPYLVNLL PCLTRTSKRP EESVQETLAA AVPKIMASFG NFANDNEIKV LLKAFIANLK SSSPTIRRTA AGSAVSICQH SRRTQYFYSW LLNVLLGLLV
301  PVEDEHSTLL ILGVLLTLRY LVPLLQQQVK DTSLKGSFGV TRKEMEVSPS AEQLVQVYEL TLHHTQHQDH NVVTGALELL QQLFRTPPPE LLQTLTAVGG
401  IGQLTAAKEE SGGRSRSGSI VELIAGGGSS CSPVLSRKQK GKVLLGEEEA LEDDSESRSD VSSSALTASV KDEISGELAA SSGVSTPGSA GHDIITEQPR
501  SQHTLQADSV DLASCDLTSS ATDGDEEDIL SHSSSQVSAV PSDPAMDLND GTQASSPISD SSQTTTEGPD SAVTPSDSSE IVLDGTDNQY LGLQIGQPQD
601  EDEEATGILP DEASEAFRNS SMALQQAHLL KNMSHCRQPS DSSVDKFVLR DEATEPGDQE NKPCRIKGDI GQSTDDDSAP LVHCVRLLSA SFLLTGGKNV
701  LVPDRDVRVS VKALALSCVG AAVALHPESF FSKLYKVPLD TTEYPEEQYV SDILNYIDHG DPQVRGATAI LCGTLICSIL SRSRFHVGDW MGTIRTLTGN
801  TFSLADCIPL LRKTLKDESS VTCKLACTAV RNCVMSLCSS SYSELGLQLI IDVLTLRNSS YWLVRTELLE TLAEIDFRLV SFLEAKAENL HRGAHHYTGL
901  LKLQERVLNN VVIHLLGDED PRVRHVAASS LIRLVPKLFY KCDQGQADPV VAVARDQSSV YLKLLMHETQ PPSHFSVSTI TRIYRGYNLL PSITDVTMEN
1001 NLSRVIAAVS HELITSTTRA LTFGCCEALC LLSTAFPVCI WSLGWHCGVP PLSASDESRK SCTVGMATMI LTLLSSAWFP LDLSAHQDAL ILAGNLLAAS
1101 APKSLRSSWA SEEEANPAAT KQEEVWPALG DRALVPMVEQ LFSHLLKVIN ICAHVLDDVA PGPAIKAALP SLTNPPSLSP IRRKGKEKEP GEQASVPLSP
1201 KKGSEASAAS RQSDTSGPVT TSKSSSLGSF YHLPSYLKLH DVLKATHANY KVTLDLQNST EKFGGFLRSA LDVLSQILEL ATLQDIGKCV EEILGYLKSC
1301 FSREPMMATV CVQQLLKTLF GTNLASQFDG LSSNPSKSQG RAQRLGSSSV RPGLYHYCFM APYTHFTQAL ADASLRNMVQ AEQENDTSGW FDVLQKVSTQ
1401 LKTNLTSVTK NRADKNAIHN HIRLFEPLVI KALKQYTTTT CVQLQKQVLD LLAQLVQLRV NYCLLDSDQV FIGFVLKQFE YIEVGQFRES EAIIPNIFFF
1501 LVLLSYERYH SKQIIGIPKI IQLCDGIMAS GRKAVTHAIP ALQPIVHDLF VLRGTNKADA GKELETQKEV VVSMLLRLIQ YHQVLEMFIL VLQQCHKENE
1601 DKWKRLSRQI ADIILPMLAK QQMHIDSHEA LGVLNTLFEI LAPSSLRPVD MLLRSMFVTP NTMASVSTVQ LWISGILAIL RVLISQSTED IVLSRIQELS
1701 FSPYLISCTV INRLRDGDST STLEEHSEGK QIKNLPEETF SRFLLQLVGI LLEDIVTKQL KVEMSEQQHT FYCQELGTLL MCLIHIFKSG MFRRITAAAT
1801 RLFRSDGCGG SFYTLDSLNL RARSMITTHP ALVLLWCQIL LLVNHTDYRW WAEVQQTPKR HSLSSTKLLS PQMSGEEEDS DLAAKLGMCN REIVRRGALI
1901 LFCDYVCQNL HDSEHLTWLI VNHIQDLISL SHEPPVQDFI SAVHRNSAAS GLFIQAIQSR CENLSTPTML KKTLQCLEGI HLSQSGAVLT LYVDRLLCTP
2001 FRVLARMVDI LACRRVEMLL AANLQSSMAQ LPMEELNRIQ EYLQSSGLAQ RHQRLYSLLD RFRLSTMQDS LSPSPPVSSH PLDGDGHVSL ETVSPDKDWY
2101 VHLVKSQCWT RSDSALLEGA ELVNRIPAED MNAFMMNSEF NLSLLAPCLS LGMSEISGGQ KSALFEAARE VTLARVSGTV QQLPAVHHVF QPELPAEPAA
2201 YWSKLNDLFG DAALYQSLPT LARALAQYLV VVSKLPSHLH LPPEKEKDIV KFVVATLEAL SWHLIHEQIP LSLDLQAGLD CCCLALQLPG LWSVVSSTEF
2301 VTHACSLIYC VHFILEAVAV QPGEQLLSPE RRTNTPKAIS EEEEEVDPNT QNPKYITAAC EMVAEMVESL QSVLALGHKR NSGVPAFLTP LLRNIIISLA
2401 RLPLVNSYTR VPPLVWKLGW SPKPGGDFGT AFPEIPVEFL QEKEVFKEFI YRINTLGWTS RTQFEETWAT LLGVLVTQPL VMEQEESPPE EDTERTQINV
2501 LAVQAITSLV LSAMTVPVAG NPAVSCLEQQ PRNKPLKALD TRFGRKLSII RGIVEQEIQA MVSKRENIAT HHLYQAWDPV PSLSPATTGA LISHEKLLLQ
2601 INPERELGSM SYKLGQVSIH SVWLGNSITP LREEEWDEEE EEEADAPAPS SPPTSPVNSR KHRAGVDIHS CSQFLLELYS RWILPSSSAR RTPAILISEV
2701 VRSLLVVSDL FTERNQFELM YVTLTELRRV HPSEDEILAQ YLVPATCKAA AVLGMDKAVA EPVSRLLEST LRSSHLPSRV GALHGVLYVL ECDLLDDTAK
2801 QLIPVISDYL LSNLKGIAHC VNIHSQQHVL VMCATAFYLI ENYPLDVGPE FSASIIQMCG VMLSGSEEST PSIIYHCALR GLERLLLSEQ LSRLDAESLV
2901 KLSVDRVNVH SPHRAMAALG LMLTCMYTGK EKVSPGRTSD PNPAAPDSES VIVAMERVSV LFDRIRKGFP CEARVVARIL PQFLDDFFPP QDIMNKVIGE
3001 FLSNQQPYPQ FMATVVYKVF QTLHSTGQSS MVRDWVMLSL SNFTQRAPVA MATWSLSCFF VSASTSPWVA AILPHVISRM GKLEQVDVNL FCLVATDFYR
3101 HQIEEELDRR AFQSVLEVVA APGSPYHRLL TCLRNVHKVT TC
```

B)

```
1    MATLEKLMKA FESLKSFQQQ QQQQQQQQQQ QQQQQQQQPP PPPPPPPPPQ LPQPPPQAQP LLPQPQPPPP PPPPPPGPAV AEEPLHRPKK ELSATKKDRV
101  NHCLTICENI VAQSVRNSPE FQKLLGIAME LFLLCSDDAE SDVRMVADEC LNKVIKALMD SNLPRLQLEL YKEIKKNGAP RSLRAALWRF AELAHLVRPQ
201  KCRPYLVNLL PCLTRTSKRP EESVQETLAA AVPKIMASFG NFANDNEIKV LLKAFIANLK SSSPTIRRTA AGSAVSICQH SRRTQYFYSW LLNVLLGLLV
301  PVEDEHSTLL ILGVLLTLRY LVPLLQQQVK DTSLKGSFGV TRKEMEVSPS AEQLVQVYEL TLHHTQHQDH NVVTGALELL QQLFRTPPPE LLQTLTAVGG
401  IGQLTAAKEE SGGRSRSGSI VELIAGGGSS CSPVLSRKQK GKVLLGEEEA LEDDSESRSD VSSSALTASV KDEISGELAA SSGVSTPGSA GHDIITEQPR
501  SQHTLQADSV DLASCDLTSS ATDGDEEDIL SHSSSQVSAV PSDPAMDLND GTQASSPISD SSQTTTEGPD SAVTPSDSSE IVLDGTDNQY LGLQIGQPQD
601  EDEEATGILP DEASEAFRNS SMALQQAHLL KNMSHCRQPS DSSVDKFVLR DEATEPGDQE NKPCRIKGDI GQSTDDDSAP LVHCVRLLSA SFLLTGGKNV
701  LVPDRDVRVS VKALALSCVG AAVALHPESF FSKLYKVPLD TTEYPEEQYV SDILNYIDHG DPQVRGATAI LCGTLICSIL SRSRFHVGDW MGTIRTLTGN
801  TFSLADCIPL LRKTLKDESS VTCKLACTAV RNCVMSLCSS SYSELGLQLI IDVLTLRNSS YWLVRTELLE TLAEIDFRLV SFLEAKAENL HRGAHHYTGL
901  LKLQERVLNN VVIHLLGDED PRVRHVAASS LIRLVPKLFY KCDQGQADPV VAVARDQSSV YLKLLMHETQ PPSHFSVSTI TRIYRGYNLL PSITDVTMEN
1001 NLSRVIAAVS HELITSTTRA LTFGCCEALC LLSTAFPVCI WSLGWHCGVP PLSASDESRK SCTVGMATMI LTLLSSAWFP LDLSAHQDAL ILAGNLLAAS
1101 APKSLRSSWA SEEEANPAAT KQEEVWPALG DRALVPMVEQ LFSHLLKVIN ICAHVLDDVA PGPAIKAALP SLTNPPSLSP IRRKGKEKEP GEQASVPLSP
1201 KKGSEASAAS RQSDTSGPVT TSKSSSLGSF YHLPSYLKLH DVLKATHANY KVTLDLQNST EKFGGFLRSA LDVLSQILEL ATLQDIGKCV EEILGYLKSC
1301 FSREPMMATV CVQQLLKTLF GTNLASQFDG LSSNPSKSQG RAQRLGSSSV RPGLYHYCFM APYTHFTQAL ADASLRNMVQ AEQENDTSGW FDVLQKVSTQ
1401 LKTNLTSVTK NRADKNAIHN HIRLFEPLVI KALKQYTTTT CVQLQKQVLD LLAQLVQLRV NYCLLDSDQV FIGFVLKQFE YIEVGQFRES EAIIPNIFFF
1501 LVLLSYERYH SKQIIGIPKI IQLCDGIMAS GRKAVTHAIP ALQPIVHDLF VLRGTNKADA GKELETQKEV VVSMLLRLIQ YHQVLEMFIL VLQQCHKENE
1601 DKWKRLSRQI ADIILPMLAK QQMHIDSHEA LGVLNTLFEI LAPSSLRPVD MLLRSMFVTP NTMASVSTVQ LWISGILAIL RVLISQSTED IVLSRIQELS
1701 FSPYLISCTV INRLRDGDST STLEEHSEGK QIKNLPEETF SRFLLQLVGI LLEDIVTKQL KVEMSEQQHT FYCQELGTLL MCLIHIFKSG MFRRITAAAT
1801 RLFRSDGCGG SFYTLDSLNL RARSMITTHP ALVLLWCQIL LLVNHTDYRW WAEVQQTPKR HSLSSTKLLS PQMSGEEEDS DLAAKLGMCN REIVRRGALI
1901 LFCDYVCQNL HDSEHLTWLI VNHIQDLISL SHEPPVQDFI SAVHRNSAAS GLFIQAIQSR CENLSTPTML KKTLQCLEGI HLSQSGAVLT LYVDRLLCTP
2001 FRVLARMVDI LACRRVEMLL AANLQSSMAQ LPMEELNRIQ EYLQSSGLAQ RHQRLYSLLD RFRLSTMQDS LSPSPPVSSH PLDGDGHVSL ETVSPDKDWY
2101 VHLVKSQCWT RSDSALLEGA ELVNRIPAED MNAFMMNSEF NLSLLAPCLS LGMSEISGGQ KSALFEAARE VTLARVSGTV QQLPAVHHVF QPELPAEPAA
2201 YWSKLNDLFG DAALYQSLPT LARALAQYLV VVSKLPSHLH LPPEKEKDIV KFVVATLEAL SWHLIHEQIP LSLDLQAGLD CCCLALQLPG LWSVVSSTEF
2301 VTHACSLIYC VHFILEAVAV QPGEQLLSPE RRTNTPKAIS EEEEEVDPNT QNPKYITAAC EMVAEMVESL QSVLALGHKR NSGVPAFLTP LLRNIIISLA
2401 RLPLVNSYTR VPPLVWKLGW SPKPGGDFGT AFPEIPVEFL QEKEVFKEFI YRINTLGWTS RTQFEETWAT LLGVLVTQPL VMEQEESPPE EDTERTQINV
2501 LAVQAITSLV LSAMTVPVAG NPAVSCLEQQ PRNKPLKALD TRFGRKLSII RGIVEQEIQA MVSKRENIAT HHLYQAWDPV PSLSPATTGA LISHEKLLLQ
2601 INPERELGSM SYKLGQVSIH SVWLGNSITP LREEEWDEEE EEEADAPAPS SPPTSPVNSR KHRAGVDIHS CSQFLLELYS RWILPSSSAR RTPAILISEV
2701 VRSLLVVSDL FTERNQFELM YVTLTELRRV HPSEDEILAQ YLVPATCKAA AVLGMDKAVA EPVSRLLEST LRSSHLPSRV GALHGVLYVL ECDLLDDTAK
2801 QLIPVISDYL LSNLKGIAHC VNIHSQQHVL VMCATAFYLI ENYPLDVGPE FSASIIQMCG VMLSGSEEST PSIIYHCALR GLERLLLSEQ LSRLDAESLV
2901 KLSVDRVNVH SPHRAMAALG LMLTCMYTGK EKVSPGRTSD PNPAAPDSES VIVAMERVSV LFDRIRKGFP CEARVVARIL PQFLDDFFPP QDIMNKVIGE
3001 FLSNQQPYPQ FMATVVYKVF QTLHSTGQSS MVRDWVMLSL SNFTQRAPVA MATWSLSCFF VSASTSPWVA AILPHVISRM GKLEQVDVNL FCLVATDFYR
3101 HQIEEELDRR AFQSVLEVVA APGSPYHRLL TCLRNVHKVT TC
```

**Figure S3. Primary sequence coverage maps of EXPI293F HTT$^{1\text{-}3144}$ Q23 digested with trypsin.** A) 77% coverage obtained from solution, and B) 62% coverage from a gel band. NB: table displays HTT Q21 sequence as UniProt database used in sequence search.

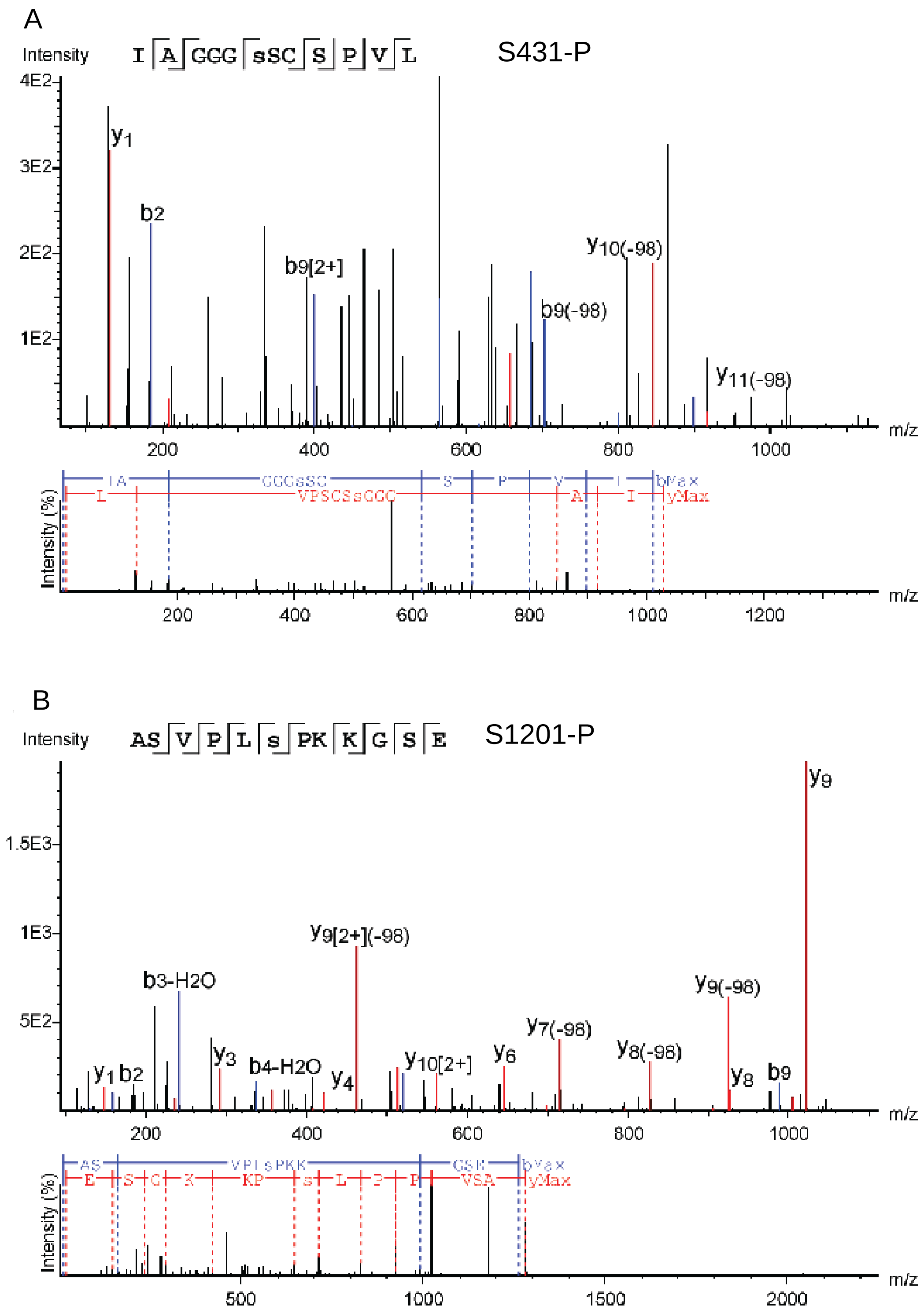


**Figure S4. Exemplary spectra of pepsin digested HTT$^{1\text{-}3144}$ Q23 from Sf9.** Full data can be found through PRIDE (2) with accession PXD010865.

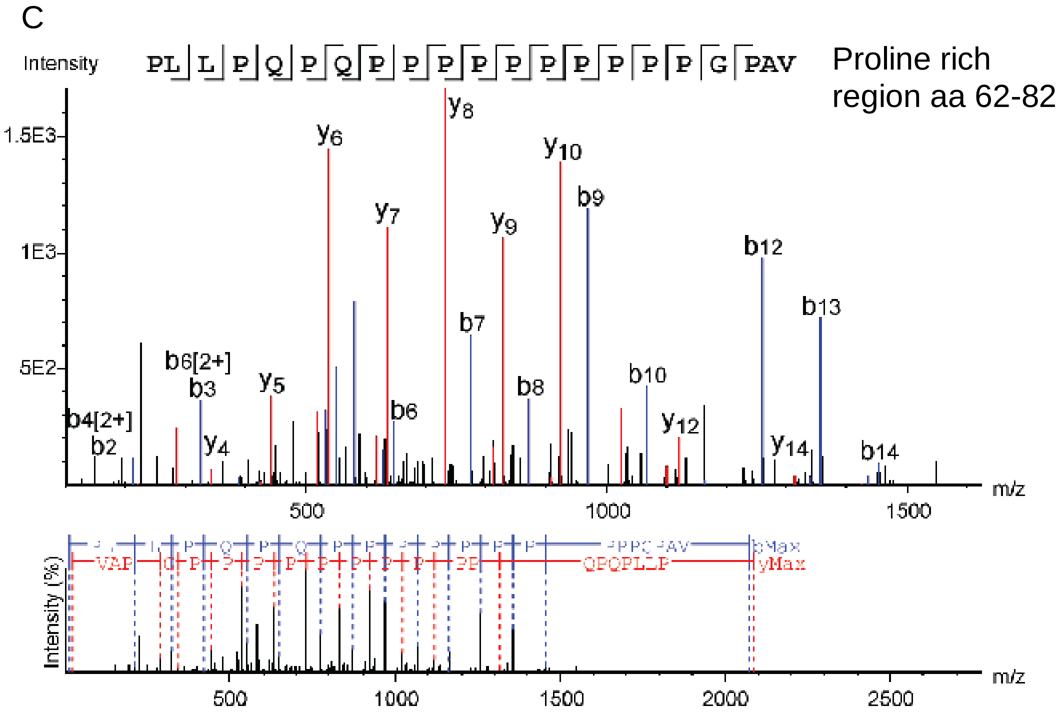


**Figure S4 continued.**

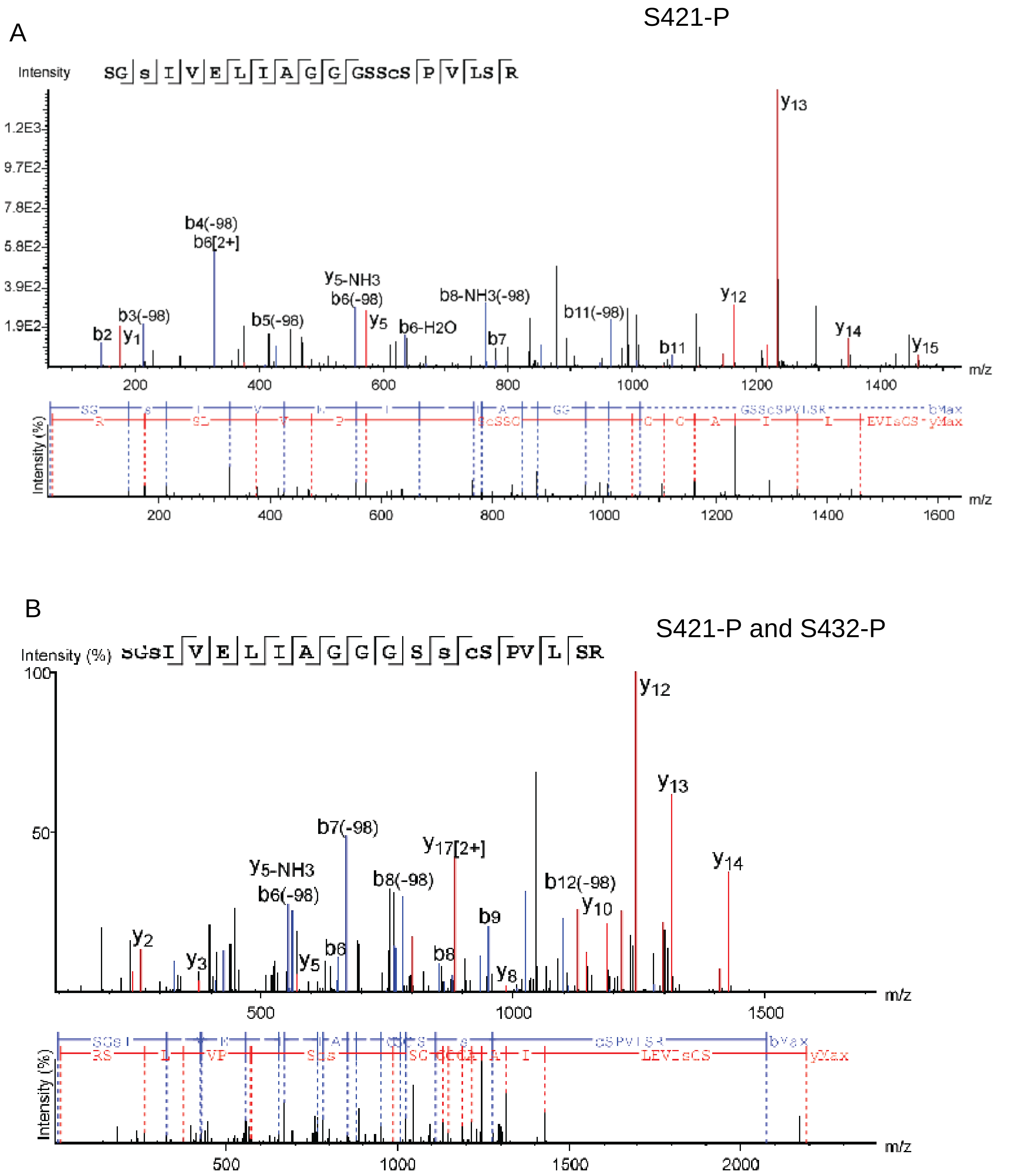


**Figure S5. Exemplary spectra of trypsin or lysargiNase digested HTT$^{1\text{-}3144}$ Q23 from Sf9.** Full data can be found through PRIDE (2) with accession PXD010865.

C

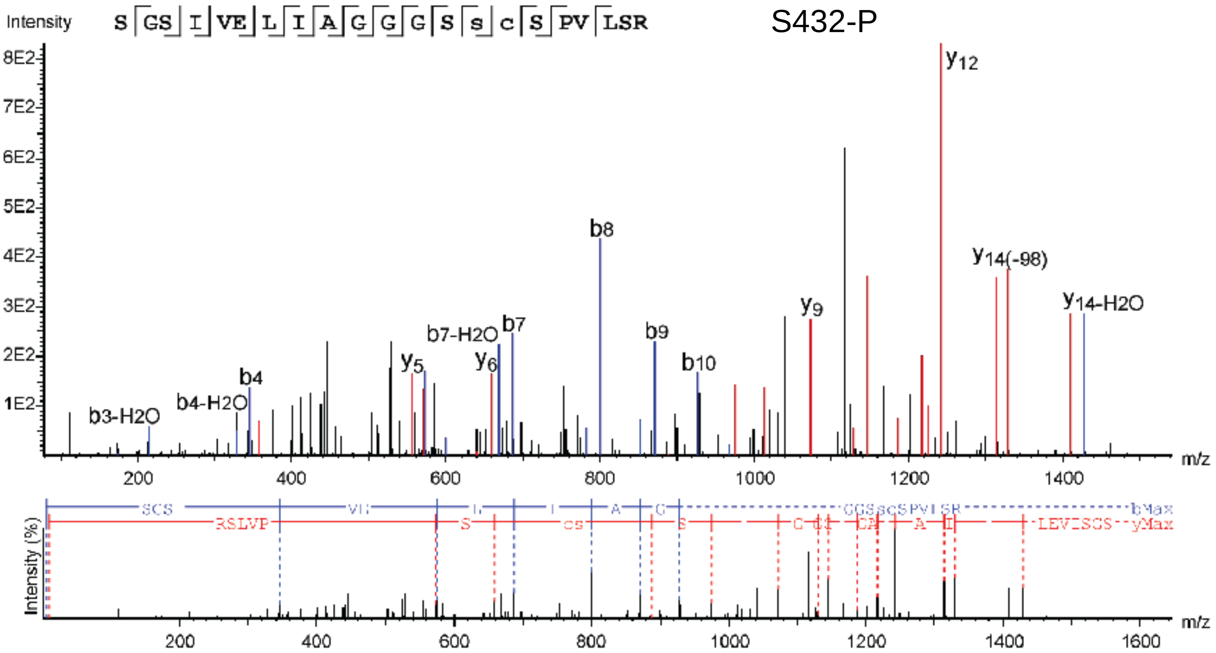


D

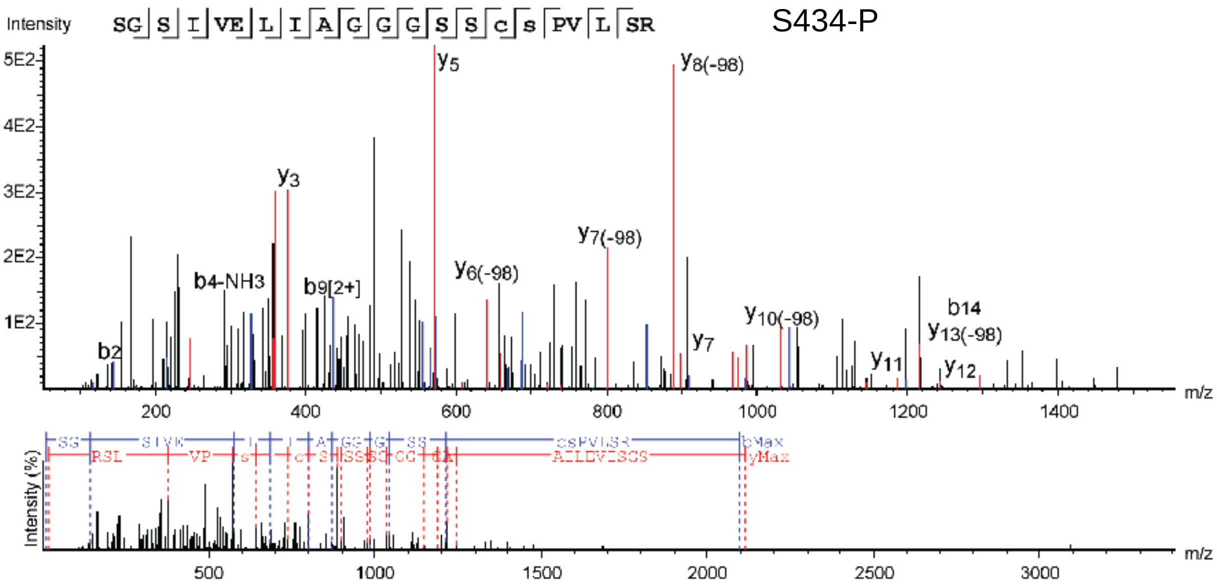


**Figure S5 continued.**

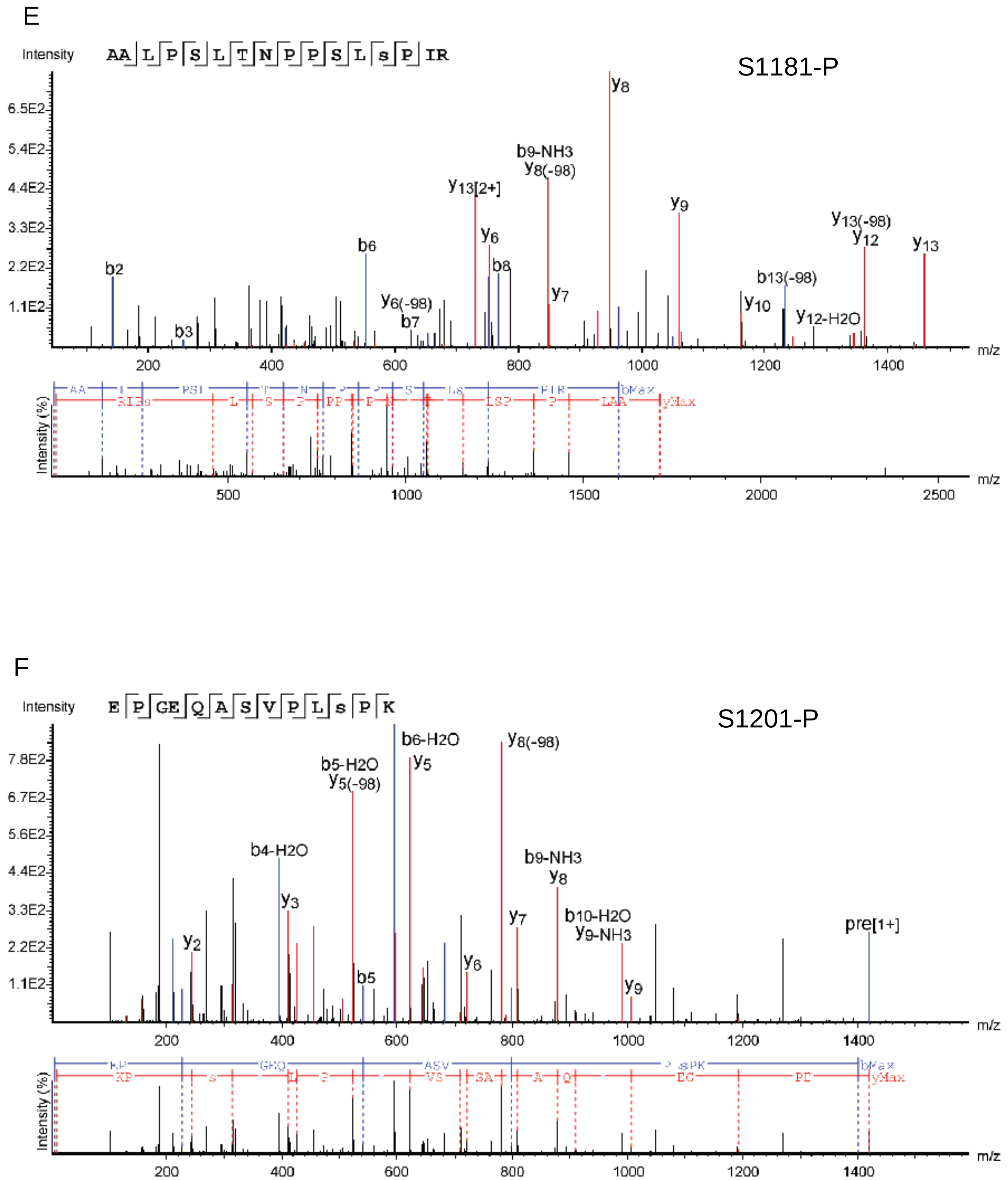


**Figure S5 continued.**

G

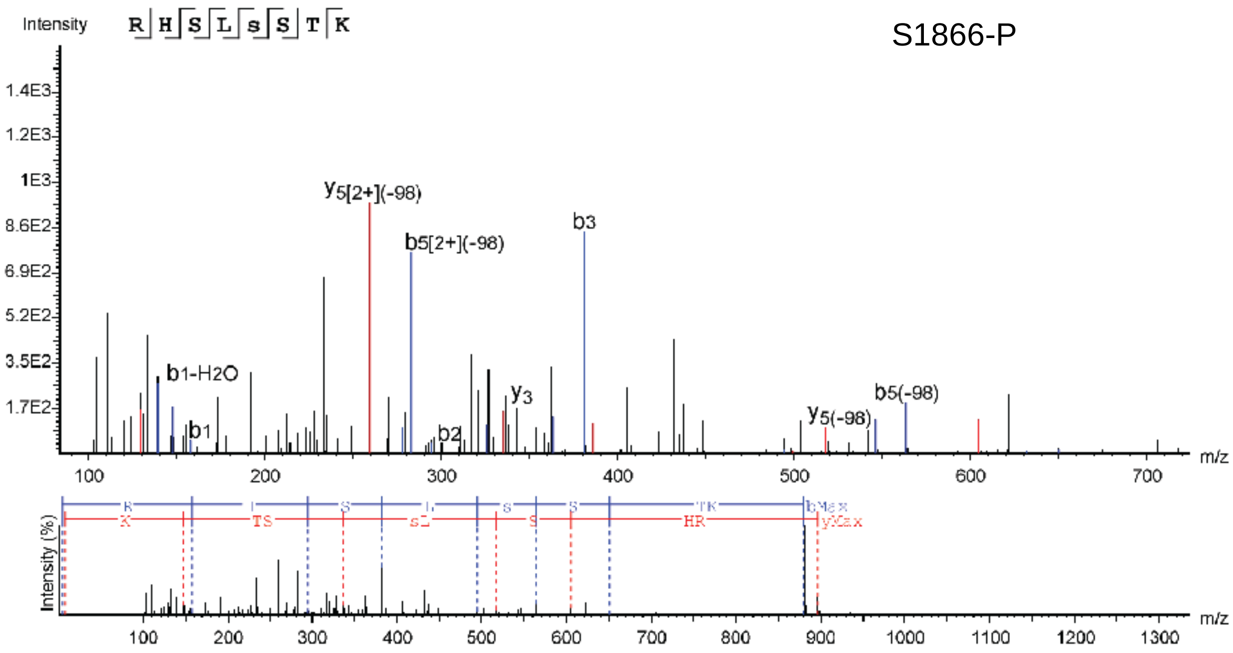


H

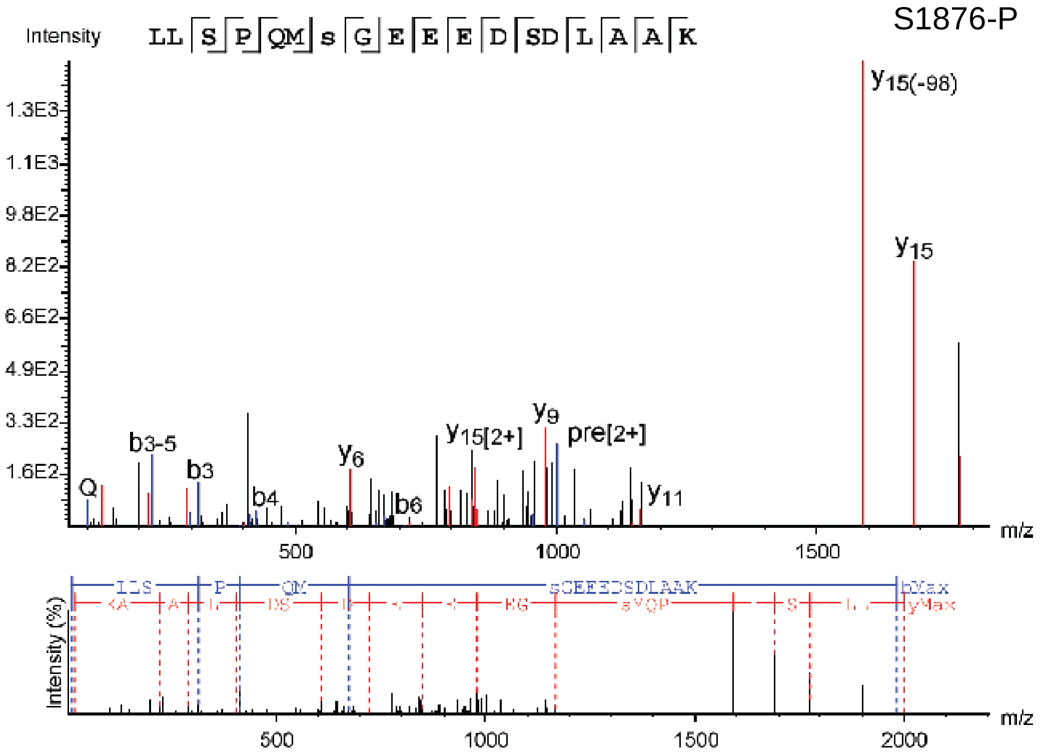


**Figure S5 continued.**

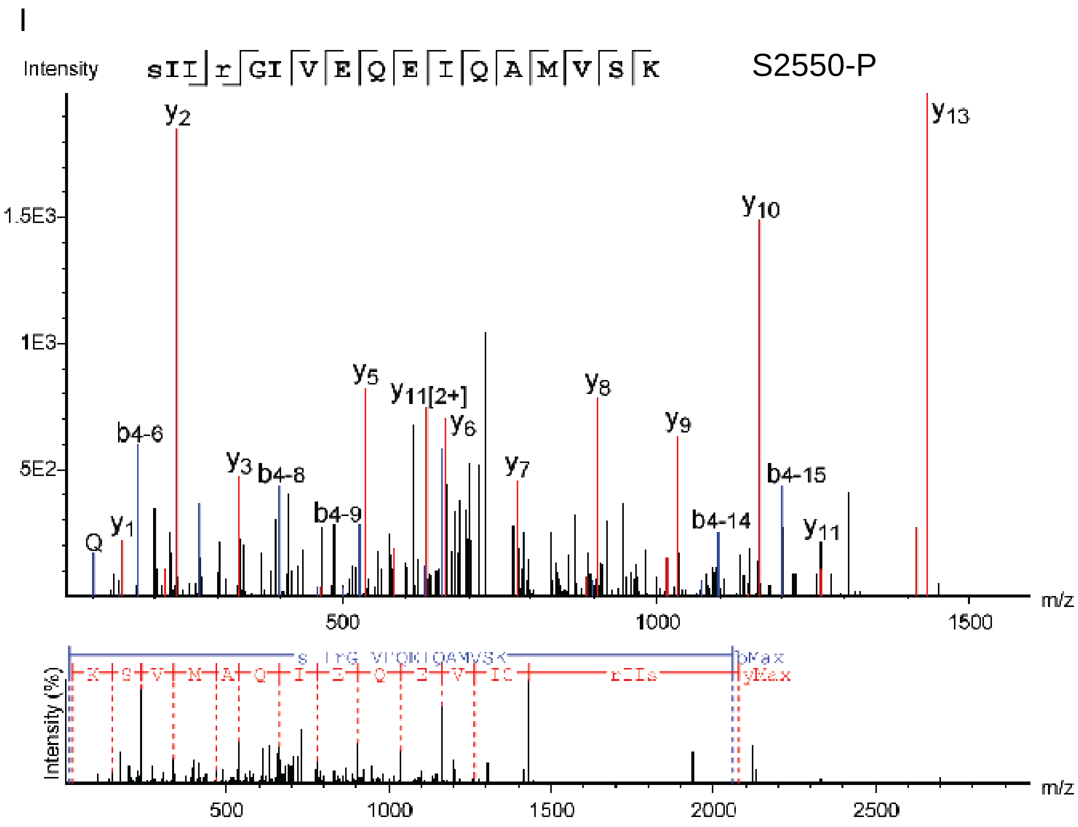


**Figure S5 continued.**

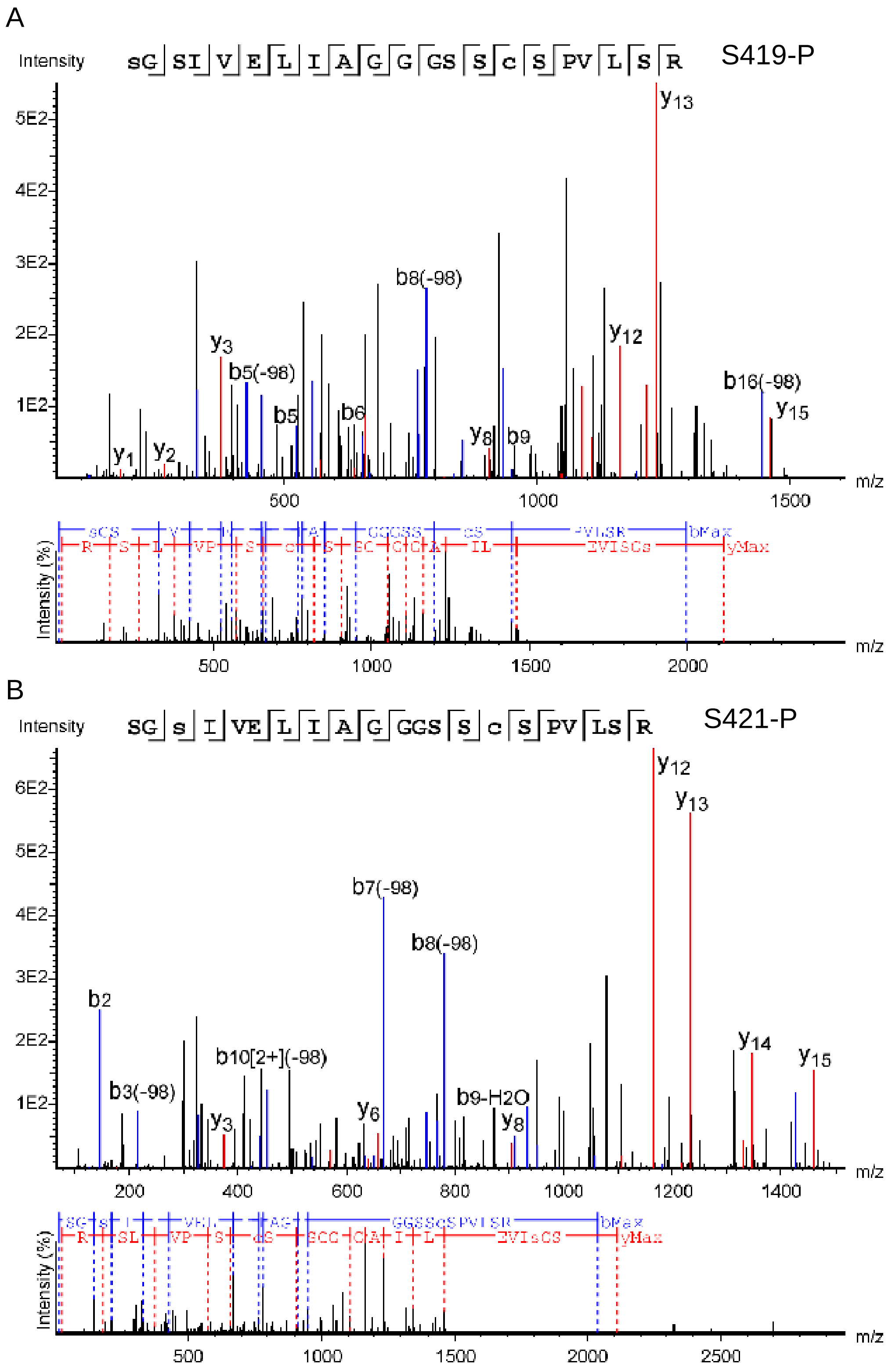


**Figure S6. Exemplary spectra of trypsin digested HTT$^{1-3144}$ Q23 from EXPI293F.**

Full data can be found through PRIDE (2) with accession PXD010865.

C

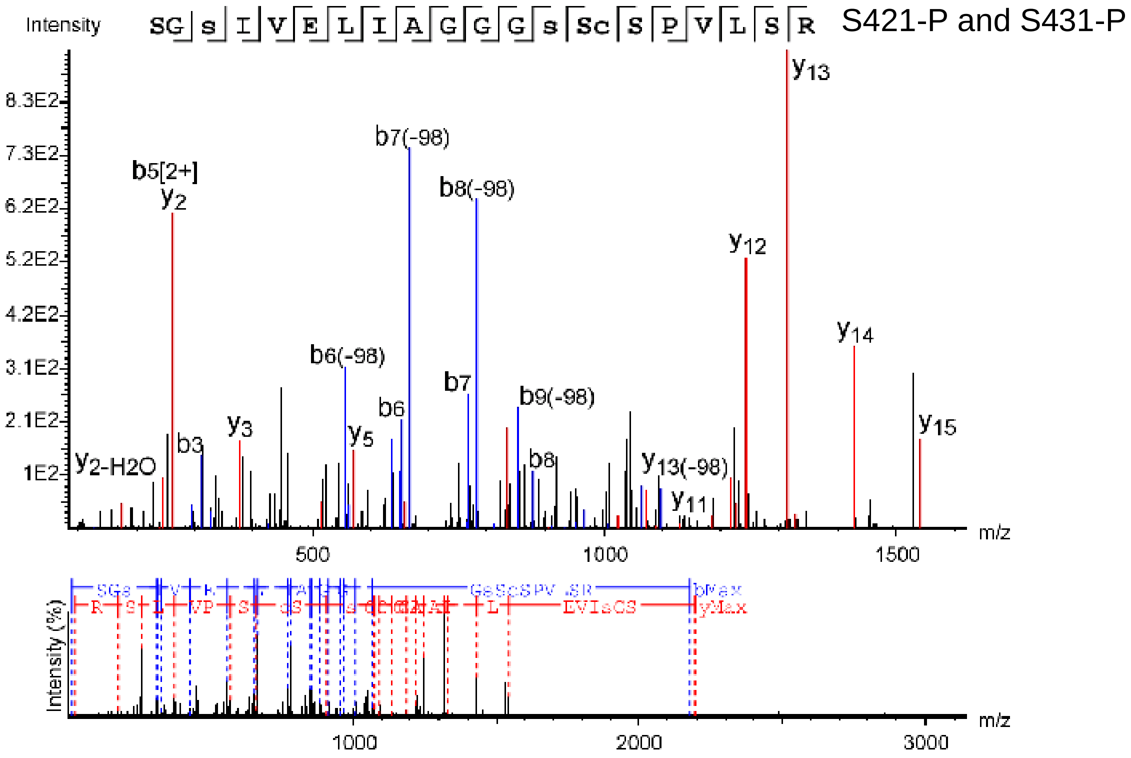


D

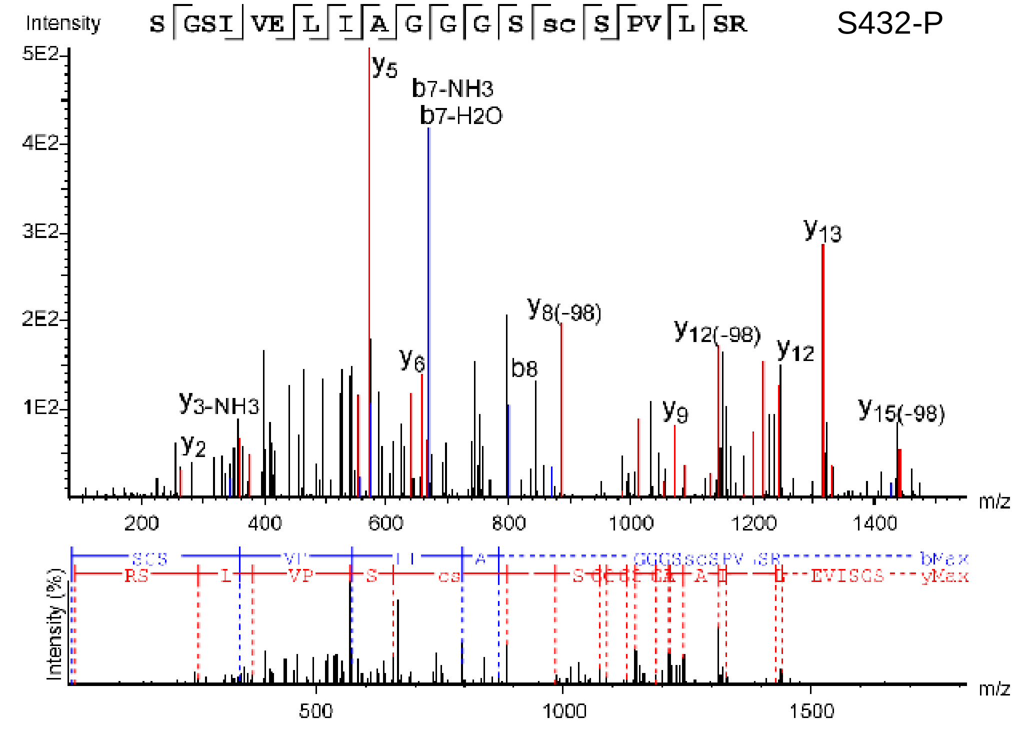


**Figure S6 continued.**

E

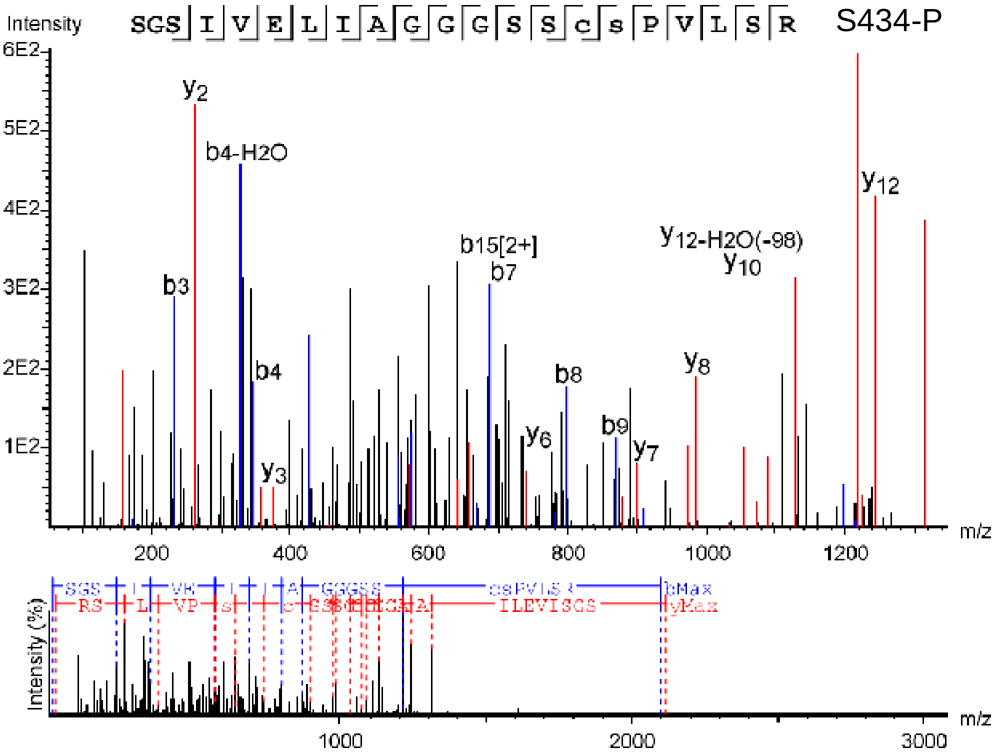


F

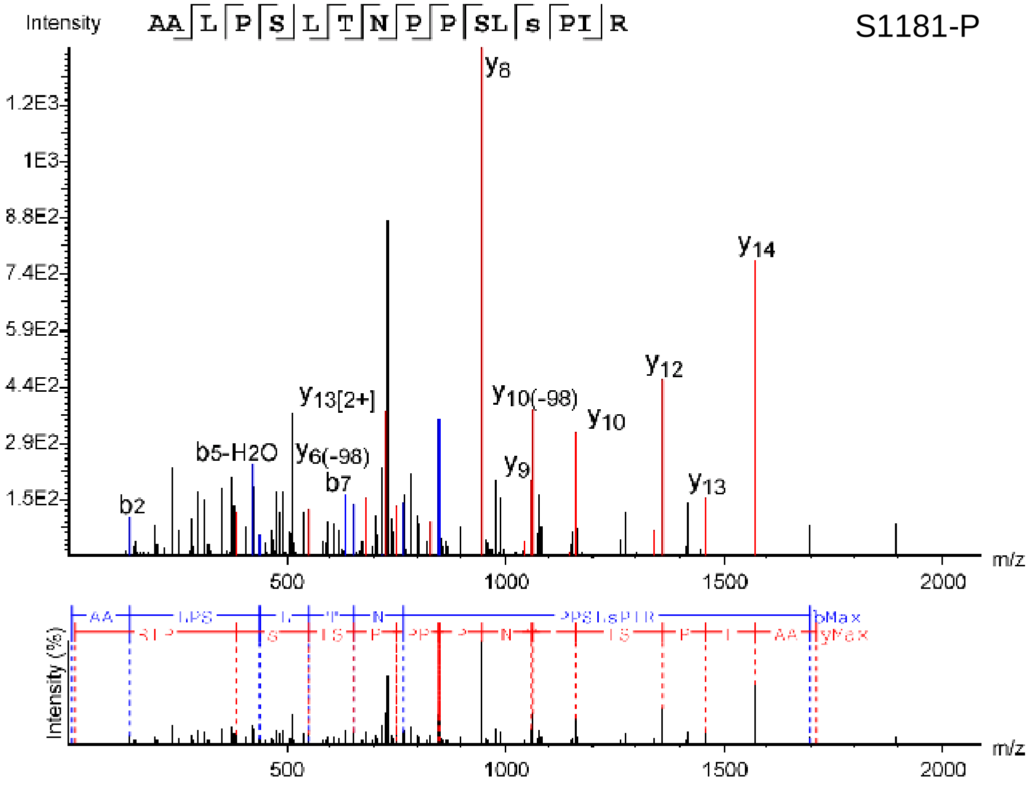


**Figure S6 continued.**

G

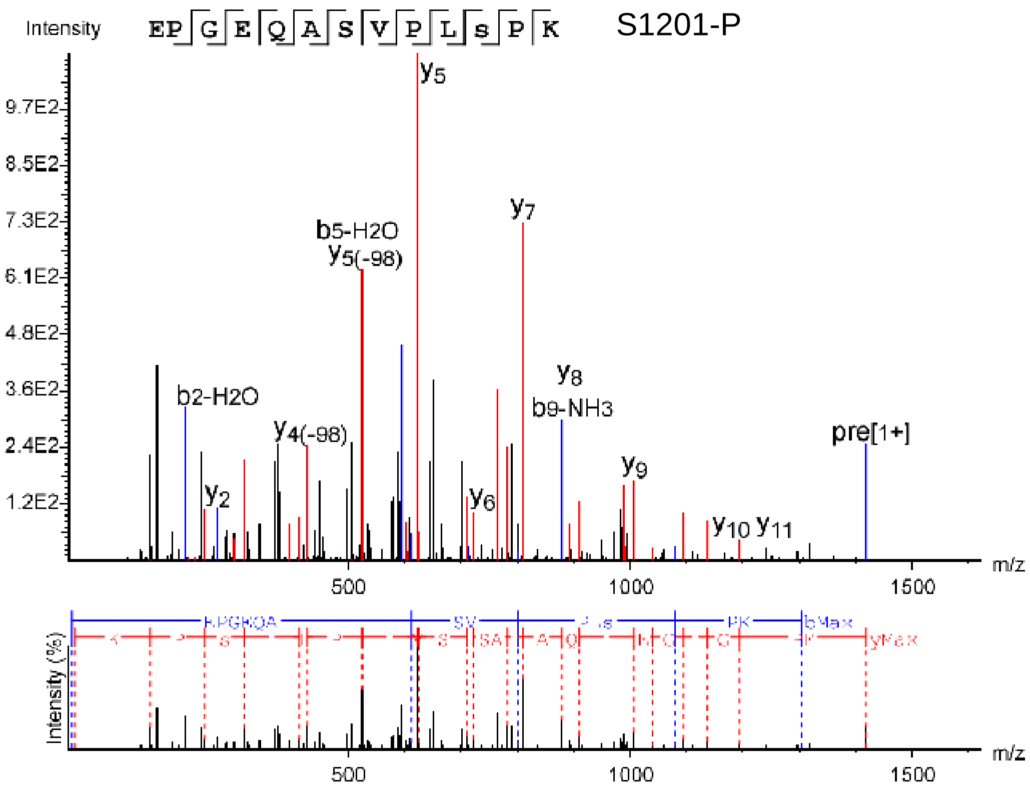


H

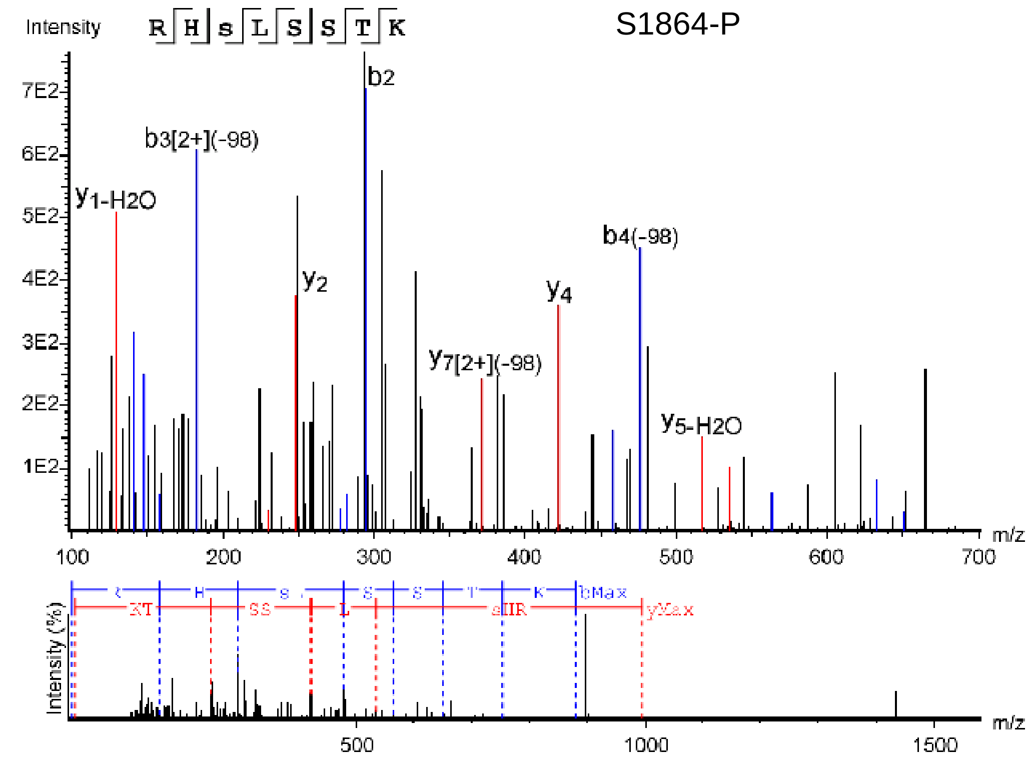


**Figure S6 continued.**

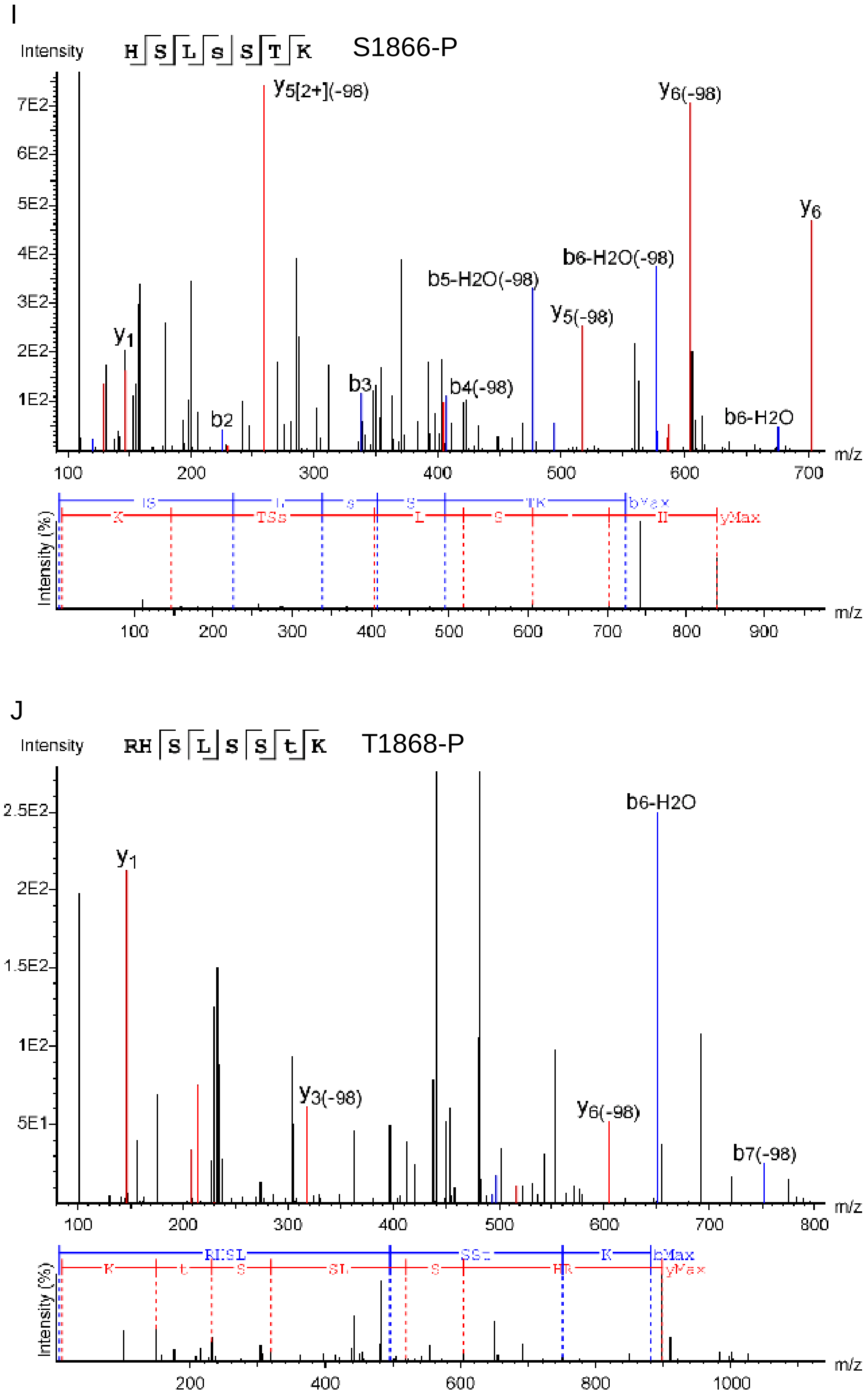


**Figure S6 continued.**

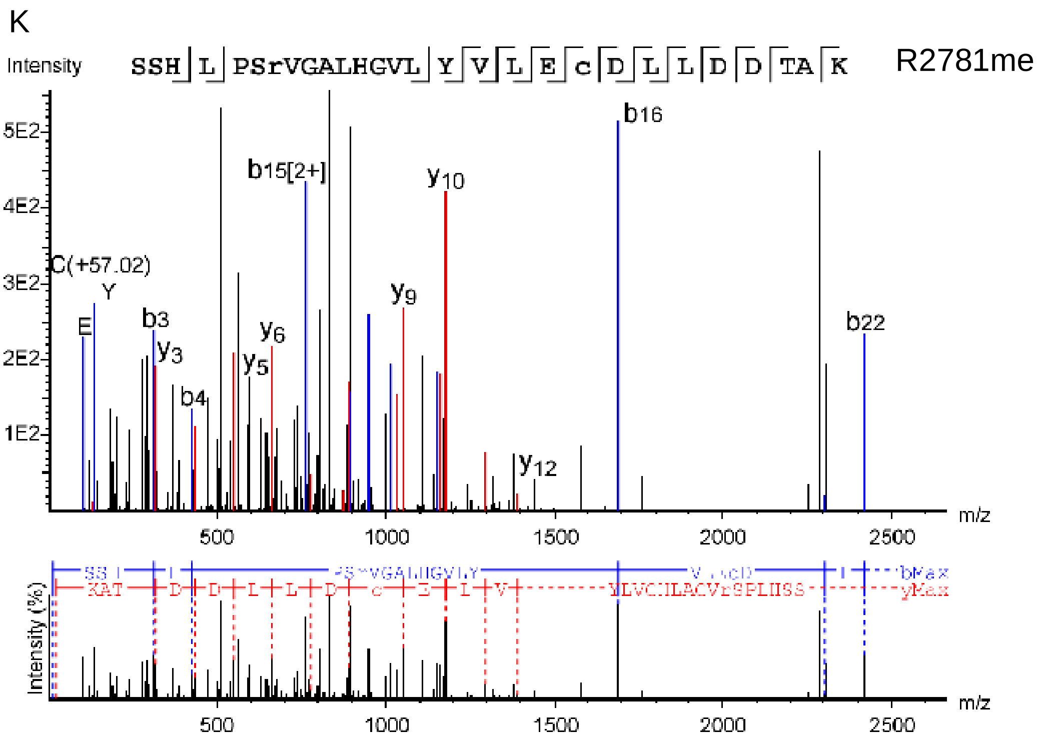


**Figure S6 continued.**

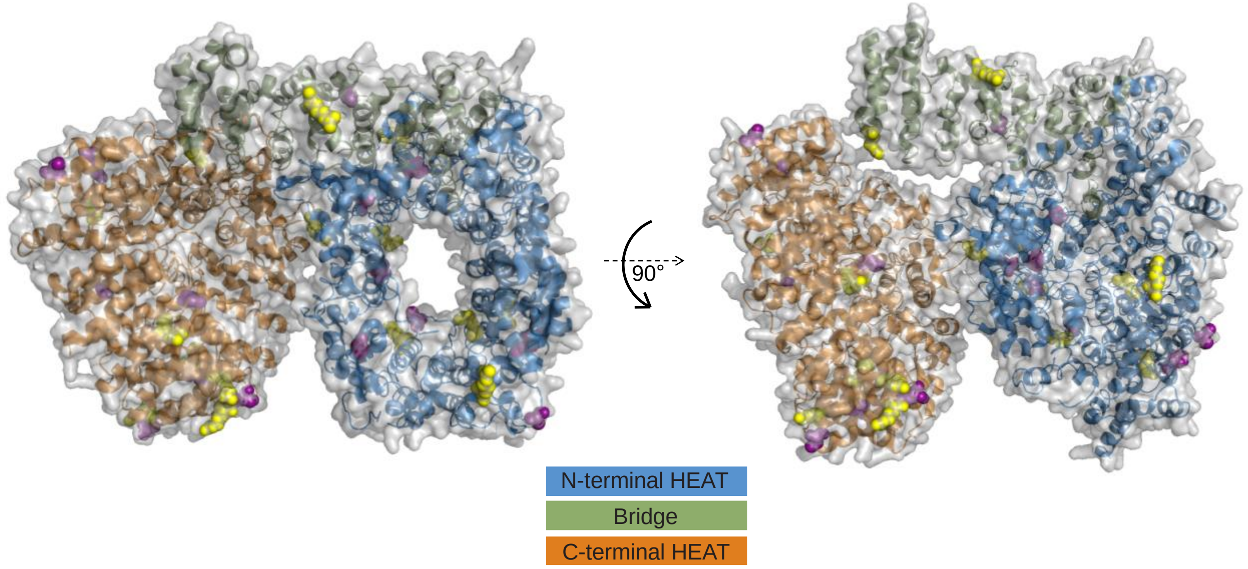


**Figure S7. Mapping HTT posttranslational modifications identified from HTT$^{1\text{-}3144}$ samples from Sf9 and EXPI293F cells onto the HTT structure.**
HTT is viewed top-down looking through the void in the N-terminal HEAT domain on the right hand side. Phosphorylation sites are shown in pink and all other modification sites are shown in green. As these samples were expressed in the absence of the stabilising HAP40 protein, it is likely that the more conformationally flexible apo HTT protein molecule would have greater exposure of different domain surfaces that would permit more sites to be modified than might be estimated from assessing the HTT-HAP40 molecule.
